# Differential interferon-α subtype immune signatures suppress SARS-CoV-2 infection

**DOI:** 10.1101/2021.05.20.444757

**Authors:** J. Schuhenn, T.L. Meister, D. Todt, T. Bracht, K. Schork, J.-N. Billaud, C. Elsner, N Heinen, Z. Karakoese, S. Haid, S. Kumar, L. Brunotte, M. Eisenacher, J. Chen, Z Yuan, T. Pietschmann, B. Wiegmann, H. Beckert, C. Taube, VTK. Le-Trilling, M. Trilling, A. Krawczyk, S. Ludwig, B. Sitek, E. Steinmann, U. Dittmer, K. Sutter, S. Pfaender

**Author notes:** Equally contributing author (J. Schuhenn, T.L. Meister, K. Sutter, S. Pfaender).

## Abstract

Type I interferons (IFN-I) exert pleiotropic biological effects during viral infections, balancing virus control versus immune-mediated pathologies and have been successfully employed for the treatment of viral diseases. Humans express twelve IFN-alpha (α) subtypes, which activate downstream signalling cascades and result in distinct patterns of immune responses and differential antiviral responses. Inborn errors in type I IFN immunity and the presence of anti-IFN autoantibodies account for very severe courses of COVID-19, therefore, early administration of type I IFNs may be protective against life-threatening disease. Here we comprehensively analysed the antiviral activity of all IFNα subtypes against SARS-CoV-2 to identify the underlying immune signatures and explore their therapeutic potential. Prophylaxis of primary human airway epithelial cells (hAEC) with different IFNα subtypes during SARS-CoV-2 infection uncovered distinct functional classes with high, intermediate and low antiviral IFNs. In particular IFNα5 showed superior antiviral activity against SARS-CoV-2 infection. Dose-dependency studies further displayed additive effects upon co-administered with the broad antiviral drug remdesivir in cell culture. Transcriptomics of IFN-treated hAEC revealed different transcriptional signatures, uncovering distinct, intersecting and prototypical genes of individual IFNα subtypes. Global proteomic analyses systematically assessed the abundance of specific antiviral key effector molecules which are involved in type I IFN signalling pathways, negative regulation of viral processes and immune effector processes for the potent antiviral IFNα5. Taken together, our data provide a systemic, multi-modular definition of antiviral host responses mediated by defined type I IFNs. This knowledge shall support the development of novel therapeutic approaches against SARS-CoV-2.

## Main

Without the capacity to produce or recognize interferons (IFN), mammalian hosts rapidly succumb in case of viral infections. Accordingly, humans with loss-of-function mutations in the IFN signalling pathway even fail to control attenuated viruses. Therefore., IFNs are indispensable mediators of the first immediate intrinsic cellular defences against invading pathogens, such as viruses. So far, three different types of IFNs, types I, II and III, have been identified and classified based on their genetic, structural, and functional characteristics as well as receptor usages^1–3^. Type I IFNs are among the first line of antiviral defence due to the ubiquitous expression of the surface receptor IFNAR consisting of two subunits IFNAR1 and IFNAR2. In humans, the type I IFN family comprises IFNβ, IFNε, IFNκ, IFNω and twelve IFNα subtypes. The latter code for the distinct human IFNα proteins: IFNα1, −2, −4, −5, −6, −7, −8, −10, −14, −16, −17 and −21, encoded by 14 nonallelic genes including one pseudogene and two genes that encode identical proteins (IFNα13 and IFNα1). The overall identity of the IFNα proteins ranges from 75 to 99% amino acid sequence identity^1,4^. Despite their binding to the same cellular receptor, their antiviral and antiproliferative potencies differ considerably^5–10^. As a general event in terms of signal transduction, IFNα subtypes engage the IFNAR1/2 receptor and initiate a signal transduction cascade resulting in the phosphorylation of receptor-associated janus tyrosine kinases culminating in downstream signalling events including the activation of IFN-stimulated gene factor 3 (ISGF3) consisting of phosphorylated STAT1 and STAT2 and the IFN regulatory factor 9. ISGF3 binding to the IFN-stimulated response elements (ISRE), in promotor regions of various genes, initiates the transcriptional activation of a large number of IFN-stimulated genes (ISGs), which elicit direct antiviral, anti-proliferative and immunoregulatory properties^11^. It is largely elusive, why different IFNα proteins exhibit distinct effector functions. Different receptor affinities and/or interaction interfaces within the IFNAR have been discussed which may account for the observed variability in the biological activity^12,13^. Furthermore, the dose, the cell type, the timing and the present cytokine milieu might further affect the IFN effector response^14^. In the absence of specific antiviral drugs, treatment of patients with type I IFNs is often considered as first therapeutic response, given its successful clinical application against viral infections^15,16^. Recently, type III IFNs (IFN-lambda, IFNλ) received significant attention and are currently explored in clinical trials^17^. IFNλ binds to the type III IFN receptor, which is preferentially expressed on epithelial cells and certain myeloid cells^18^, resulting in restricted cell signalling and compartmentalized activity. Especially at epithelial surface barriers, IFNλ mount an effective local innate immune response, by conferring viral control and inducing immunity without generating systemic activation of the immune system which could trigger pathologic inflammatory responses. Signal transduction cascades of type I and type III IFNs are considered to be rather similar resulting in overlapping ISG signatures, however, type I IFN signalling leads to a more rapid induction and decline of ISG expression^19^.

The outbreak of novel viruses, as exemplified by the recent emergence of *Severe Acute Respiratory Syndrome Coronavirus-2* (SARS-CoV-2), causing the disease COVID-19 has emphasised the urgent need for fast and effective therapeutic strategies. Indeed, type I IFN treatment is currently explored as emergency treatment against COVID-19 in various clinical trials^20–22^, and it was already shown that SARS-CoV-2 is sensitive to type I IFNs^23^ and ISGs^24^. Given their large genome size, CoVs have evolved a variety of strategies circumventing the host innate immune reaction, including evasion strategies targeting type I IFN signalling^23,25–27^. Along those lines, recent studies showed significantly decreased interferon activity in COVID-19 patients who developed more severe disease^28^, highlighting the importance of IFN in controlling viral infection. Against viruses, pegylated IFNα2 is approved and frequently administered in clinical settings. However, common side effects include the occurrence of flu-like symptoms, haematological toxicity, elevated transaminases, nausea, fatigue, and psychiatric sequelae, which often result from systemic activation of the immune system^29^. Given the described distinct biological properties of IFNα subtypes, we comprehensively studied their antiviral effect against SARS-CoV-2 in comparison to another respiratory virus (influenza A virus), and we aimed to explore SARS-CoV-2-specific immune signatures that could contribute to an efficient viral clearance. Accordingly, the aim of this study was two-fold: I) to identify underlying immune-signatures crucial for controlling SARS-CoV-2 infection and II) to explore the therapeutic potential of IFNα subtypes in SARS-CoV-2 infection.

## Results

### IFNα subtypes differentially inhibit SARS-CoV-2

In order to determine the antiviral potencies of the twelve different IFNα subtypes against SARS-CoV-2, we pre-treated VeroE6 cells with two doses (1000 units per mL (U/mL) and 100 U/mL). We included IFNλ3 (1000 ng/mL and 100 ng/mL), since its potent antiviral activity against SARS-CoV-2 and other respiratory pathogens has been documented^30,31^. Following treatment for 16 hours, cells were subsequently infected with SARS-CoV-2 and viral replication was quantified by determining infectious viruses (TCID_50_/mL) and genome amplification. Interestingly, we observed a differential antiviral pattern for the individual subtypes, with IFNα5, α4, α14 and IFNλ3 exhibiting the strongest antiviral effects with up to 10^5^ fold reduction in viral titres (Figure 1A and Extended Data Figure 1A). Immunofluorescence analysis of VeroE6 cells pre-treated with IFNα5, IFNα7 and IFNα16 confirmed their different antiviral activities against SARS-CoV-2 (Figure 1B). To determine the inhibitory concentration 50 (IC_50_), we performed dose-response analyses covering concentrations from 19 to 80,000 U/mL for the pre-treatment. SARS-CoV-2 replication was assessed by quantification of viral titres (TCID_50_/mL) and viral antigens applying a previously described in-cell ELISA^32^ (Table 1 and Extended Data 1B-D). Corroborating previous results, a striking clustering of the antiviral subtypes according to their antiviral potency was observed, which allowed their separation into classes of low (IC_50_ >5000 U/mL), intermediate (IC_50_: 2000-5000 U/mL) and high (IC_50_: <2000 U/mL) antiviral activities against SARS-CoV-2 (Fig. 1C-F, Extended Data 1B-D, Table 1).

**Fig. 1:**
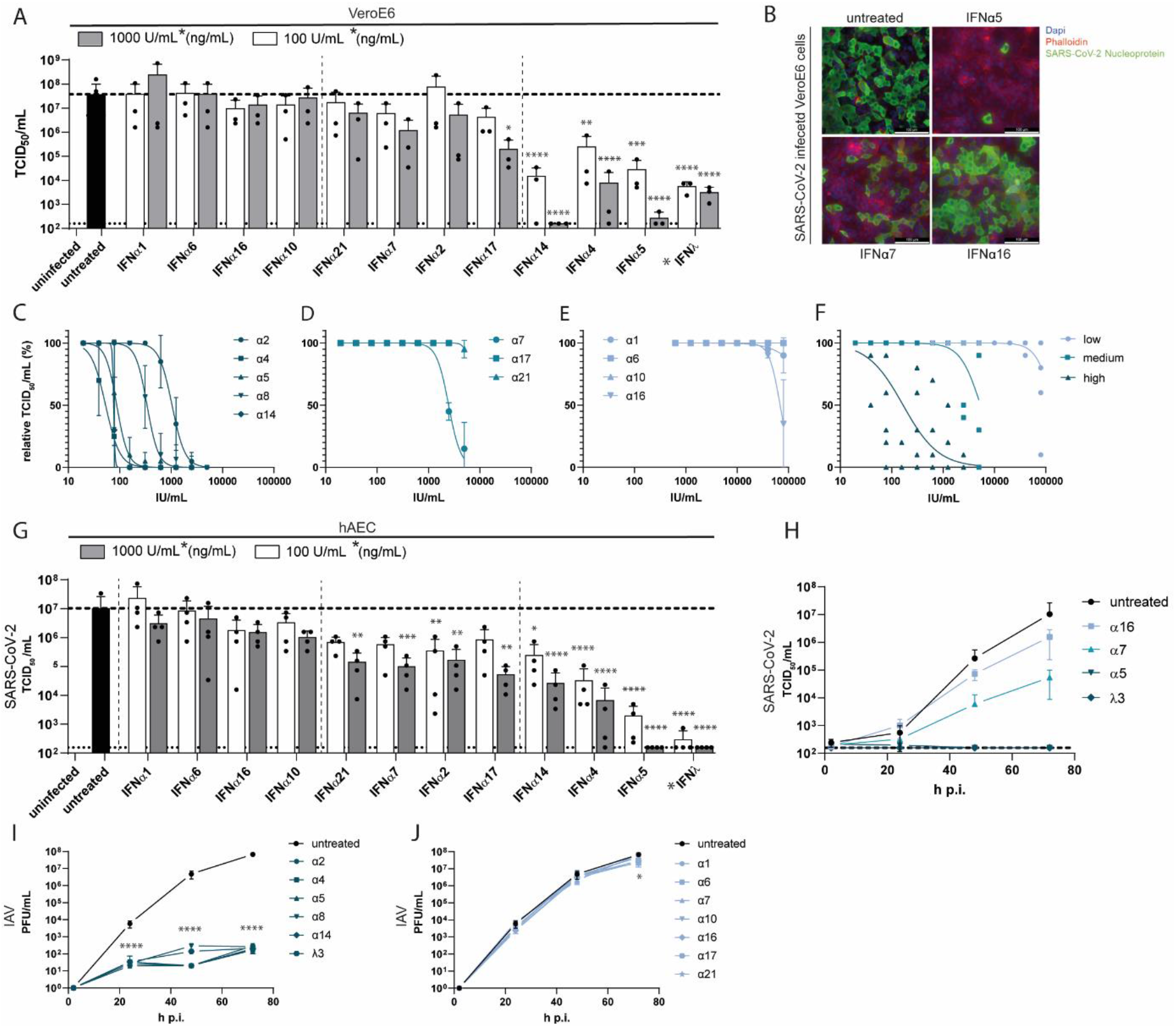
Treatment with IFNα subtypes reveals distinct antiviral effects against SARS-CoV-2. (A) Antiviral activity of IFNα subtypes (100 or 1000 U/mL) and IFNλ3 (100 or 1000 ng/mL) against SARS-CoV-2 on VeroE6 cells (TCID_50_/mL). (B) Representative immunofluorescence staining of IFN-treated SARS-CoV-2 infected VeroE6 cells. IFNα subtypes were titrated against SARS-CoV-2 on VeroE6 cells by TCID_50_ assay and the IFNs were grouped in high (C), medium (D) and low (E) antiviral pattern and the mean values of each group are plotted in (F). Antiviral activity of IFNα subtypes and IFNλ3 in SARS-CoV-2-infected primary hAECs at 72 h p.i. (G) and kinetics of four selected IFNs (H). (L-N) Antiviral activity of IFNα subtypes and IFNλ3 in Influenza A/PR8-infected primary hAECs at different timepoints post infection. Mean values of high (I) and low/not (J) antiviral IFNs are shown. (A, C-F; I, J) Mean values ± SEM are shown for n=3. (G, H) n=4. A: 100 U/mL (ng/mL): ** p=0.0035 (IFNα4); *** p=0.0002 (IFNα5); **** p<0.0001 (IFNα14, IFNλ3); 1000 U/mL (ng/mL): * p=0.0180 (IFNα17); **** p<0.0001 (IFNα4, α5, α14, λ3). G: 100 U/mL (ng/mL): * p=0.0352 (IFNα14); ** p=0.0063 (IFNα2) *** p=0.0002 (IFNα4); **** p<0.0001 (IFNα5, IFNλ3); 1000 U/mL (ng/mL): ** p=0.0028 (IFNα2) p=0.0016 (IFNα17) p=0.0021 (IFNα21) *** p=0.0003 (IFNα7); **** p<0.0001 (IFNα4, α5, α14, λ3); I: **** p<0.0001 (all IFNs, all time points J. 72hpi * p=0.0468 (IFNα16)

**Table 1:** IC_50_ values of IFNα subtypes on VeroE6 cells obtained from endpoint dilution assay.

| IFNα subtype | IC<sub>50</sub> [U/mL] |
| --- | --- |
| IFNα4 | 56.91 |
| IFNα14 | 70.73 |
| IFNα5 | 79.73 |
| IFNα8 | 327.0 |
| IFNα2 | 1026 |
| IFNα7 | 2431 |
| IFNα21 | 4944 |
| IFNα16 | >5000 |
| IFNα1 | >5000 |
| IFNα17 | >5000 |
| IFNα6 | >5000 |
| IFNα10 | >5000 |

Since VeroE6 cells are derived from African green monkey, expressing the non-human primate instead of human IFN receptor and also lack the capacity to produce IFN-I in a natural feed-forward loop^33^, we further analysed genuine target cells of SARS-CoV-2. We utilized well-differentiated primary human airway epithelial cells (hAEC), which closely resemble the *in vivo* physiology of the respiratory system, and differentiate into various cells types, resulting in ciliary movement and production of mucus ^34,35^. After IFN pre-treatment and subsequent infection with SARS-CoV-2, apical washes were monitored concerning viral replication kinetics at 33°C ^36^. Cells were lysed at 72 h post infection (p.i.) and viral progeny (Fig. 1G, H) as well as viral *M* and *N* gene expression (Extended Data 1 E-J) were determined. Again, a distinct antiviral pattern became evident (Figure 1G) defining IFN clusters of high (IFNα5, −4, −14, − IFNλ3), moderate (IFNα17, −2, −7, −21) and low antiviral activities (FNα10, −16, −6, −1) (Fig. 1H and Extended Data 1 G, J). Prototypical ISG expression patterns, as analysed by qRT-PCR, revealed subtype-specific gene expression signatures (Extended Data Figure 2A-E. In order to address if the observed antiviral activities were SARS-CoV-2-specific, we additionally tested influenza A virus (IAV/PR8) in hAECs. Interestingly, pre-treatment of hAECs with the IFN-subtypes revealed differences compared to SARS-CoV-2. In general, antiviral responses could be clustered into strong for α2, −4, −5, −8, −14 and IFNλ3 (Fig. 1I) and weak antiviral activities for IFNα1, −6, −7, −10, −16, −17 and 21 (Fig. 1J). Amongst the strong antiviral responses, we observed additional transient differences at 48 h p.i., with IFNα2, −4, −5 and −14 being slightly superior to IFNα8 and −λ3 (Fig. 1I). These results clearly demonstrate that different IFNα subtypes mediate distinct biological and temporal activities.

### IFN subtype-specific gene expression signatures

Since we observed clear differences in the biological activities of different IFNα subtypes against SARS-CoV-2, we next aimed to identify their underlying immune signatures and mechanisms. To this end, primary hAECs were pre-treated with the respective IFNs and 16 h post stimulation cellular RNA was sequenced on an Illumina NovaSeq 6000 and differentially expressed genes were sent to Ingenuity Pathway Analysis (IPA; Qiagen) for biological analysis. In order to investigate cellular responses following viral infection, we included SARS-CoV-2-infected hAECs (18 h p.i.) in our analysis. Global transcriptomic analysis revealed unique differentially expressed genes (DEGs), both up- and downregulated upon IFN-treatment ^37,38^ for each IFN (Extended Data Figure 3A) compared to mock-treated cells. Similar to the observed antiviral effects, a general clustering was apparent which showed similar expression patterns for low to intermediate antiviral subtypes (IFNα1, −6, −7, −16, −10, −21) and intermediate to high antiviral subtypes (IFNα2, −17, −14, −4, −5, −λ3). Interestingly, we observed a clear difference in the numbers of significantly up- and down-regulated genes after treatment with IFNα subtypes compared to mock-treated cells, which positively correlated with antiviral activity (Extended Data Figure 3B). Gene ontology (GO) pathway analysis revealed higher expression of genes mostly involved in antiviral immune response amongst the medium and high antiviral subtypes, as well as pathways which can be associated with protein localization, translation, oxidative phosphorylation, RNA metabolism, ER stress, signalling pathways and lymphocyte activation (Figure 2A). Strikingly, different IFNα subtypes displayed unique GO patterns with IFNα17, in contrast to other subtypes, regulating genes involved in translation, whereas the treatment with IFNα5 resulted in the strongest regulation of genes associated to signalling pathways and lymphocyte activation among all IFNs (Figure 2A). We next focussed on genes associated with antiviral responses (Figure 2B). A separation based on antiviral activity could be discerned with weak antiviral IFNα subtypes (IFNα1, −6, −16, −10) exhibiting comparatively lower expression values of specific ISGs, whereas medium to strong antiviral IFNα subtypes induced higher expression (Figure 2B). We observed two clusters that differed between low and intermediate to high IFN subtypes, with ISG15, IFI27, MX1 and others showing generally lower expression values in the low antiviral IFN subtypes. Even more pronounced were expression changes of IFIT2, IFIT1 and MX2 and others which resulted in a down-regulation for the low- and an upregulation for the intermediate to high antiviral IFN subtypes. As we aimed at identifying immune signatures that correlate with the antiviral activity against SARS-CoV-2 infection, we next evaluated DEGs with respect to distinct, intersecting and common genes amongst and between subtypes (Extended Data Figure 4A). We identified several differentially expressed genes for each subtype, with IFNα5 expressing most unique genes (1018 DEGs), followed by IFN λ3 (670 DEGs) (Figure 2C, Extended Data Figure 4B)). A comparison between high, medium and low antiviral subtypes revealed that 19 genes were commonly differentially expressed amongst all subtypes including *Mx1* and *OAS2* (Figure 2D). The most striking differences could be observed for *MX1* and *OAS2*, which expression levels clearly separated high, intermediate and low antiviral IFN subtypes (Figure 2D). Interestingly, 42 genes were differentially regulated in the high antiviral group including *RNaseL* and genes associated with regulation of transcription, signal transduction and metabolic processes (Figure 2E), as well as long non-coding RNAs. In conclusion, we could clearly demonstrate IFN subtype-specific immune signatures that could contribute to the observed differences in antiviral activity.

**Fig. 2:**
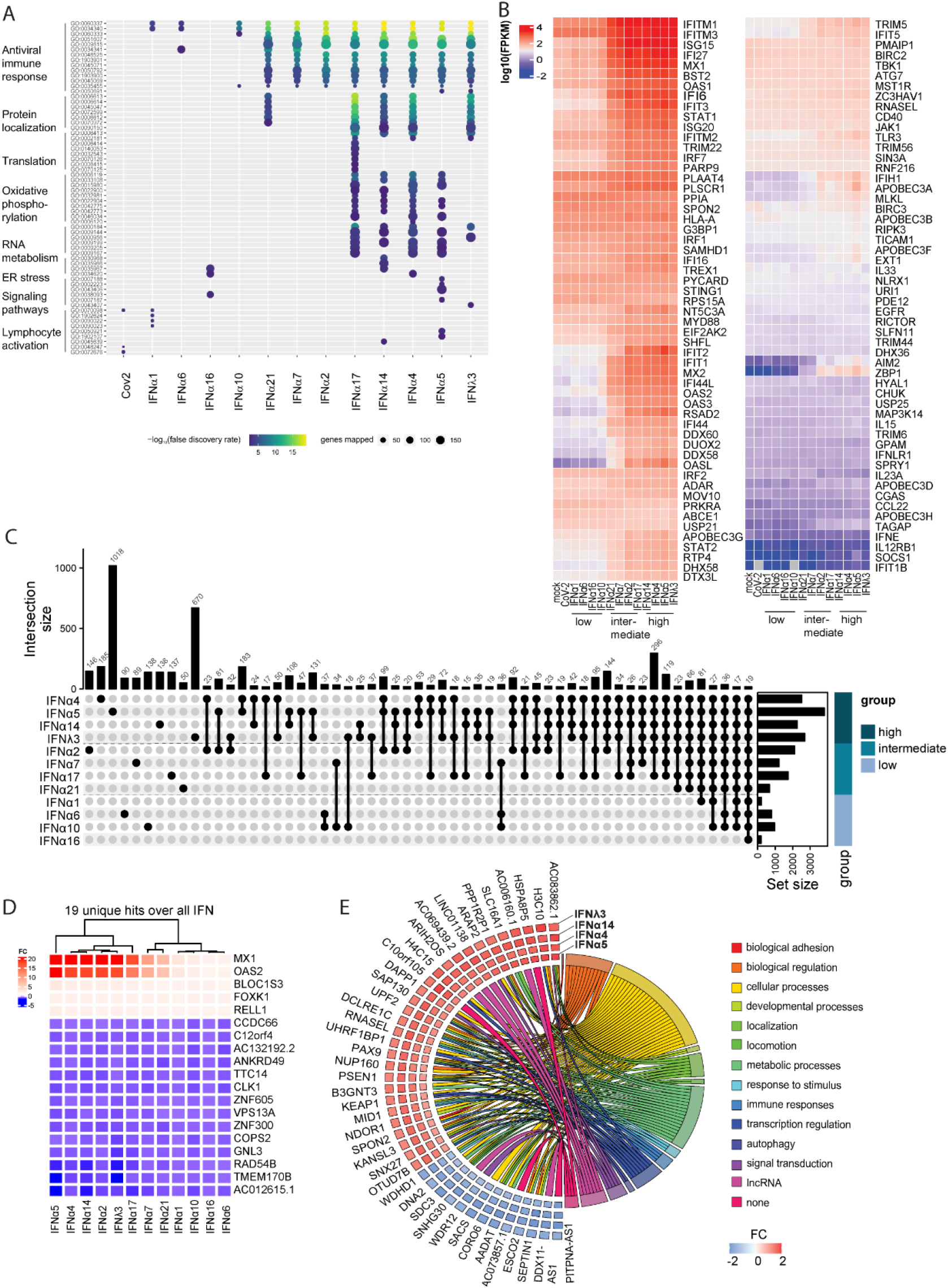
Transcriptomic analysis display IFN subtype specific immune signatures. (A-E) Transcriptomic analyses of IFN-treated (16 hours post treatment; 1000U/mL or 1000ng/mL) or SARS-CoV-2-infected (18 hours post infection) hAECs. (A) Biological processes induced by IFNs or SARS-CoV-2. (B) Heat maps displaying genes contained in antiviral response. (C) UpSet plots to summarize key differentially expressed genes (DEG). Numbers of individually or group-specific DEGs are shown as bars and numbers. The bottom right horizontal bar graph labelled Set Size shows the total number of DEGs per treatment. IFNs are plotted due to their antiviral activity in 3 groups (high, medium and low). (D) Heatmap of the 19 basal DEGs expressed by all IFNs as identified in D. (E) Plot depicting fold changes (FC) of identified 42 unique genes in the group displaying high antiviral activity and association of genes to functional categories. (A-E) n=4

### Proteomic analysis highlights key cellular factors

Our transcriptomic analysis revealed IFNα subtype-specific distinct, intersecting and common expression patterns of DEGs that most likely contribute to the differential biological activity against SARS-CoV-2. To further uncover relevant cellular effector proteins for the antiviral activity against SARS-CoV-2, we additionally performed proteomic analysis on hAECs pre-treated with IFNs. Since we had observed the strongest antiviral activity for IFNα5 and IFNλ3 we decided to further investigate their specific proteomic profile in direct comparison with IFNα7, which exhibited a moderate antiviral effect, and IFNα16, displaying a weak effect against SARS-CoV-2 infection, in order to identify key antiviral pathways, crucial in controlling coronavirus infection. To this end, primary hAECs were pre-treated with selected IFNs for 16 h. In addition to the early time point (t=0 h), where we aim to identify key cellular factors that are expressed before viral infection, we included a late time point, 72 h post treatment both in the presence (t=72 h [CoV-2]) or absence of viral infection (t=72 h [mock]), to investigate potential antiviral mechanisms and potential intervention by viral effectors (Extended Data Figure 5A). Principal component analysis (PCA) revealed a clustering according to donor and/or infection and time points (Extended data Figure 5B-D, Extended Data Table 2). In addition to host cell proteins, various viral peptides were identified, which correlate to viral titres depending on the respective donor (Extended Data Table 3, Extended Data Figure 5E). For all donors, no SARS-CoV-2 peptides could be detected following treatment with IFNα5 and IFNλ3. Pre-treatment of cells with IFN subtypes resulted in up- or down-regulation of a variety of proteins compared to untreated hAECs, depending on the IFN stimulation (Extended Data Figure 6A-C). In order to perform statistical analysis, we considered proteins that were measured in minimum three of four donors, however on/off analysis (defined as full absence of a protein in one group of a pairwise comparison) revealed additional proteins which might be of interest (Extended Data Figure 6D-F, Extended Data Table 4). GO analysis of proteins differentially abundant between untreated and IFN-treated samples at each time point (untreated vs IFN) identified enrichment of antiviral immune responses for all IFNs, except IFNα16 (Figure 3A, Extended Data Figure 7A). For IFNα16, only proteins associated with lymphocyte regulation were induced, which likely do not contribute to SARS-CoV-2 restriction in cell culture but may be very important *in vivo*. At 72 h pathways belonging to proteolysis, metabolism and protein localization were additionally enriched after treatment with IFNα5 and −λ3. The most prominent upregulated proteins, associated with IFN signalling (STAT1, MX1, ISG15, ISG20, IFI35, and others) were found to be on-off regulated and present only upon treatment with IFNα5, −α7 and −λ3. Additional ISGs including IFIT3, OAS2, and IFITM3 were on-off regulated after 72 h and CoV-2 infection except for IFNα16-treatment (Figure 3B, Extended Data Figure 7B). Interestingly, the comparison of samples in the presence or absence of SARS-CoV-2 (Mock vs CoV-2) showed a striking trend towards downregulation of proteins upon CoV-2 infection. Enrichment of biological processes associated with complement activation and O-glycan processing (Figure 3C) highlighted various complement factors (e.g. CFB, C4B and C3) as well as various mucines (e.g. Muc1, Muc16) by SARS-CoV-2, independent of IFN-treatment and resulting viral titres (Figure 3D, Extended Data Figure 7C, E, Extended Data Table 5). In contrast, the strongest biological effects on antiviral immune responses after treatment with IFNα5 and −λ3, e.g. IFN signalling as well as antigen presentation, NF-κB signalling or lymphocyte regulation were not affected by viral infection. Interestingly, proteins belonging to other pathways e.g. antigen presentation by MHC class I or proteolysis, seemed to be less abundantly represented under viral infection in the IFNα5 treated samples, a phenomenon which was not as prominent after treatment with IFNλ3 (Figure 3E, Extended Data Figure 7D). STRING analysis (Figure 3F) highlighted the presence of antiviral key effector molecules (e.g. ISG20, ISG15, IFI44L, IFIT2, IFIT3, IFI35, PML, SP100), which are involved in type I IFN signalling pathways, negative regulation of viral processes and immune effector processes amongst the most potent antiviral IFNs. In conclusion, we identified a variety of antiviral cellular effector molecules that correlate with antiviral activity and controlling coronavirus infection

**Fig. 3:**
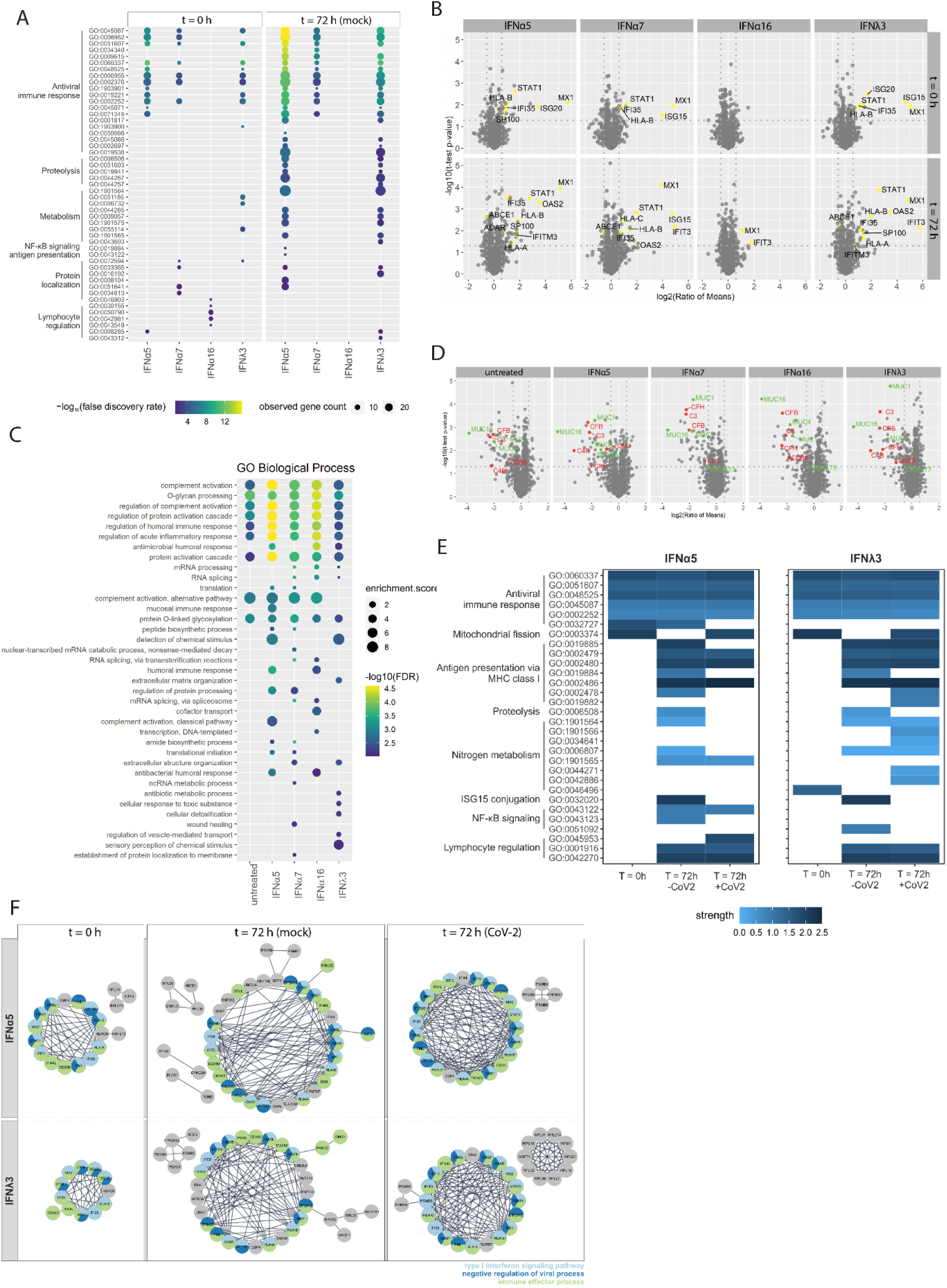
Proteomic analysis highlights key cellular mediators. (A-G) Proteomic analysis of IFN-treated (1000U/mL or 1000ng/mL) and/or SARS-CoV-2-infected hAECs. (A) Biological processes induced by IFNs 16 hours post treatment (t=0 h) or 88 hours post treatment (t=72 h). (B) Volcano plots of IFN-treated hAECs at different timepoints post treatment. Detected ISGs are coloured yellow. (C) Biological processes induced by IFNs 88 hours post treatment in the presence of SARS-CoV-2 (t=72 h) (D) Volcano plots of IFN-treated SARS-CoV-2-infected hAEC. Detected proteins are coloured due to their biological function: red = complement activation; green =O-glycan processing. (E) Heatmaps of differentially activated biological processes by highly antiviral IFNα5 and IFNλ3 compared to untreated controls at different time-points post treatment in the presence and absence of SARS-CoV-2. (F) STRING analysis of proteins increased in IFN-treated and/or SARS-CoV-2 infected hAECs and identified abundant protein-protein interactions. Proteins shown as circles and colours indicating biological processes (A-F) n=4.

### Therapeutic potential of IFNα subtypes

Currently, there are only a few approved specific antiviral drugs (e.g. monoclonal antibodies)^39,40^ for the treatment of COVID-19, which severely limit treatment options during severe clinical courses. Remdesivir, a nucleotide-analogous RNA dependent RNA Polymerase (RdRP) inhibitor originally developed as antiviral against Ebola virus, received an emergency use-approval against COVID-19 and has been employed in the clinics. Unfortunately, due to lack of evidence for recovery of critically ill patients, it is no longer recommended by the World Health Organization (WHO) as single treatment for COVID-19 ^41^). Therefore, alternative therapeutic approaches such as combination therapies are urgently needed. As we have observed the strongest antiviral effect in this study for IFNα5 we explored its therapeutic potential in comparison and in combination with remdesivir. In regard to patients viewed as an entity, prophylactic treatment with IFNs is no clinical option. Nevertheless, a treatment initiated following diagnosis can still ‘prophylactically’ condition and protect cells in the body against later infection events. To monitor the kinetics of the antiviral activity of IFNα subtypes, we treated cells either before infection (‘pre-‘) or up to 8 h post infection (‘post-‘) and studied the antiviral activity by determining viral titres as TCID_50_/mL and viral antigens by ic ELISA (Figure 4A, B). As expected, the strongest reduction in viral titres was observed upon pre-treatment with IFNα5 as cells become alerted towards an antiviral state and antiviral effectors can be transcribed or even translated prior to viral infection (Figure 4B). Intriguingly, even after viral infection was established, treatment with IFNα5 was able to significantly reduce viral titres (Figure 4B), which was also observed with the antiviral drug remdesivir (Extended Data Figure 8A). Given the clear antiviral but incomplete inhibitory effect of both treatment modalities, we next studied a potential beneficial effect of IFNα5 when co-administered with remdesivir (see Figure 4A for a schema). To this end, we analysed the antiviral effect upon pre-treatment as well as post-treatment of an established infection. To quantify the interaction between the two antiviral drugs, the observed combination response was compared to the expected effect using the Loewe additivity model, with δ-scores above 10 indicating synergistic effects. Combination therapies in VeroE6 cells revealed an additive antiviral activity, with over 90 % viral inhibition upon pre-treatment in the highest concentrations of both drugs tested and a Loewe synergistic score of 8.504 (Figure 4C, D) without any effect on cytotoxicity (Extended Data Figure 8B). Similarly, post-treatment resulted in a dose-dependent, additive viral inhibition with over 70 % (Figure 4E, F). To confirm these findings, we analysed selected combinations of IFNα5 with remdesivir post-infection in hAEC. For this we combined low doses (0.313 μM remdesivir, 0.2444 U/mL IFNα5), medium doses (0.63 μM remdesivir, 15.625 U/mL IFNα5) and high doses (2.5 μM remdesivir, 1.953 U/mL IFNα5), and observed in all combinations an additive therapeutic effect when co-administered 8 h post infection (Figure 3G-I). Taken together, we provide evidence that co-administration of direct antiviral drugs together with potent IFNα subtypes clearly impaired viral replication and might provide an alternative therapeutic approach.

**Fig. 4:**
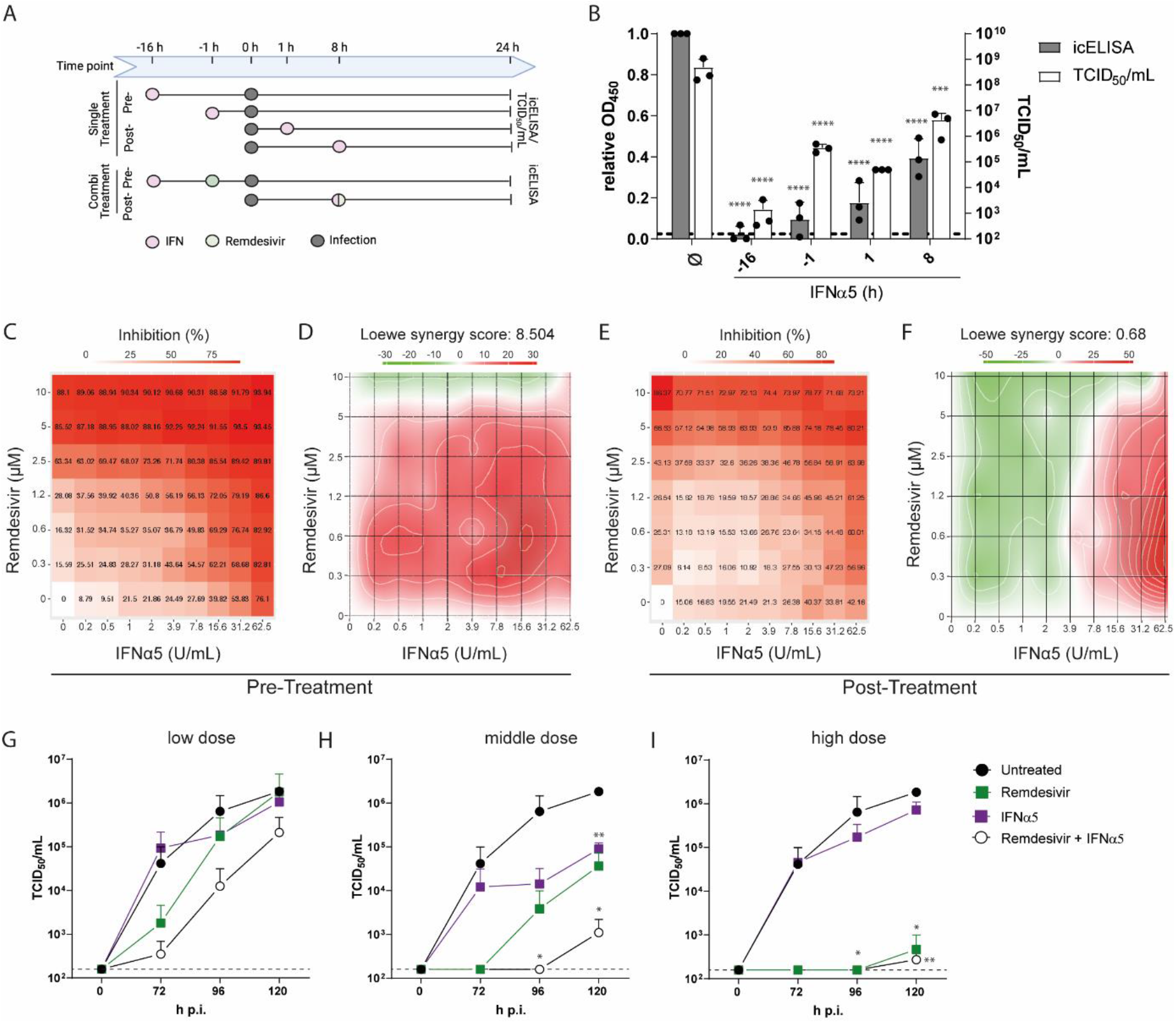
Therapeutic potential of highly antiviral IFNα subtype 5. (A-F) Single and combined treatments of IFNα5 and remdesivir in SARS-CoV-2 infected VeroE6 cells. (A) Schematic depiction of treatment. (B) Pre- and post-treatments of VeroE6 cells by icELISA (grey bars) and TCID_50_ assay (white bars). (C) Inhibition of SARS-CoV-2 infection and (D) analysis of drug combination experiments using SynergyFinder web application 16 hours before infection. (E) Inhibition of SARS-CoV-2 infection and (F) analysis of drug combination experiments using SynergyFinder web application^65^ (doi: 10.1093/bioinformatics/btx162) 8 hours post infection. (G-I) remdesivir and IFNα5 combinational treatment 8 hours post infection of hAECs with low doses (0.313 μM remdesivir, 0.2444 U/mL IFNα5; G), medium doses (0.63 μM remdesivir, 15.625 U/mL IFNα5; H) and high doses (2.5 μM remdesivir, 1.953 U/mL IFNα5; I) (B-I) n=3. B: icELISA (grey bars) **** p<0.0001; TCID50/mL (white bars) *** p=0.0003 (+8) **** p<0.0001 (−16, −1, +1) H: 96h p. i. * p=0.0205 (remdesivir + IFNα5);120h p. i. * p=0.0113 (remdesivir + IFNα5) ** p=0.0041 (IFNα5) I: 96h p. i. * p=0.0205 (remdesivir, remdesivir + IFNα5);120h p. i. ** p=0.0081 (remdesivir) ** p=0.0015 (remdesivir + IFNα5)

## Discussion

Type I interferons serve as one of the first lines of defence and are induced almost immediately upon viral encounters. Type I IFN foster intrinsic immunity, stimulate innate immunity, and recruit and orchestrate adaptive immunity. They can modulate the immune system in several ways, by exerting a wide range of biological activities including antiviral, antiproliferative, immunomodulatory and regulatory activities. Importantly, impaired type I IFN activity are correlated with severe courses of COVID-19, highlighting their clinical importance^42^. Accordingly, defectiveness to type I IFNs significantly contributes to disease severity and genetic polymorphisms decreasing IFN-I production are associated with more severe cases of COVID-19^43–45^. Furthermore, pegylated IFNα2a therapy in patients with inborn errors of type I IFN immunity prevented severe COVID-19 disease^46^. In addition to the impaired type I IFN response triggered by SARS-CoV-2, recent studies have demonstrated the development of autoantibodies that can neutralize type I IFNs^44,47^. To evade the antiviral effects of type I IFNs, viruses have evolved various strategies to suppress IFN induction. SARS-CoV-2 codes for several proteins that have been implicated in type I IFN antagonism, thereby compromising host responses and favouring viral replication^48^. Thus, early administration of IFN-I might be an effective treatment option for COVID-19 patients. The IFN-I family consists of multiple IFNα subtypes, which are highly conserved, and they all signal through the same ubiquitously expressed IFNAR1/2. Activation of various downstream signalling cascades implicates that the IFNα subtypes share some overlapping functions, but also possess unique properties. Upon pre-treatment of cells with twelve distinct IFNα subtypes, we observed cluster-specific antiviral patterns which were distinct between different viruses. These differential antiviral functions cannot be explained solely by the binding affinity to both receptor subunits as IFNα5 and IFNα4 exhibit a median affinity to IFNAR1 and IFNAR2 in the range of 0.94-3 μM and 2.1-3.8 nM, respectively^12^. Furthermore, the increased gene induction did not correlate with binding affinity to IFNAR1 or 2, as those IFNs with the highest binding affinity to IFNAR2 (IFNα10, 17, 6, 14, 7) did not induce significantly higher numbers of differentially expressed genes. In IFN-treated gut biopsies of chronically HIV-infected patients, the numbers of induced genes by different type I IFNs (IFNα1, α2, α5, α8, α14 and β) were not associated with binding affinity or ISRE activation^49^. Importantly, it has been shown that the different type I IFNs induced a specific pattern of genes, which are involved in various biological processes^49^. We observed distinct antiviral patterns, that could be clearly clustered into high, intermediate and low antiviral effects against SARS-CoV-2. Interestingly, we identified 19 genes that were common between all groups, indicative of a basal IFN response. On top of that basal response, we identified several genes that were distinct-, intersecting- or commonly differentially regulated between the high and/or medium group. Our dataset enabled us to identify expression patterns that can be correlated with antiviral activity against SARS-CoV-2. Foremost, antiviral immune responses were significantly dysregulated in the moderate and high antiviral groups. Nevertheless, several biological processes e.g. such associated with protein localization, translation or ER stress, displayed variable induction patterns depending on the IFNα subtype. Proteomic analysis confirmed expression of IFN effector molecules in high and moderate antiviral subtypes. We mostly identified factors involved in type I IFN signalling pathways, negative regulation of viral processes and immune effector processes. These results clearly demonstrate unique and overarching properties of different IFNα subtypes. Another group recently reported that saturated concentrations (1000pg/mL) of IFNα subtypes against HIV-1 *in vitro* induced similar levels of 25 canonical ISGs^50^. The authors concluded from these 25 ISGs that the overall difference between all subtypes is only quantitatively, but not qualitatively, implying that the transcription of 25 genes is fully sufficient to describe the whole interferome^51^. We similarly observe a clear difference in the magnitude of differential regulated genes, that likely contributes to the observed antiviral patterns. Nevertheless, as demonstrated with IAV, these patterns do affect virus replication to a different extent, indicating that individual IFNα subtypes might have discriminative clinical effects. Due to its known antiviral activity and its clinical administration in chronic viral infections, type I IFNs, specifically IFNα2 or IFNβ, were already used in a variety of different clinical trials in patients with mild or severe COVID-19. During SARS-CoV-2 infection, two phases can be observed: 1) an early phase with weak IFNα/β production and limited antiviral responses and 2) an excessive inflammatory immune response which can give rise to cytokine storms or acute respiratory distress syndrome. Therefore, a potential beneficial effect of IFN treatment must occur early during infection to not exacerbate hyperinflammation. Early subcutaneous administration of IFNβ in combination with lopinavir/ritonavir and ribavirin in patients with mild to moderate COVID-19 led to a significant reduction of symptoms, shortening the duration of viral shedding and hospital stay^22^. Pulmonary administration of type I IFNs might reduce systemic side effects, while increasing type I IFN concentrations in the infected epithelial cells. Inhaled or nebulized IFNα2b with arbidol or IFNβ-1b showed faster recovery from SARS-CoV-2 infection and decreased levels of inflammatory cytokines^20,21^. Furthermore, prophylactic intranasal application of IFNα2a/b in health care workers in China completely prevented new SARS-CoV-2 infections ^52^. A recent report from SARS-CoV-2 infection in golden hamsters demonstrated a systemic inflammation in distal organs like brain or intestine^53^. They hypothesized that virus-derived molecular patterns and not infectious SARS-CoV-2 were disseminated to the periphery leading to systemic inflammation and increased IFN signatures. These observations might further highlight the need to apply type I IFNs via intranasal route or inhalation, as the IFN response in the periphery is already highly stimulated and a systemic administration would not further increase the antiviral host immune response. We clearly demonstrated the additive benefit of combining treatment of type I IFN with a direct acting antiviral, e.g. remdesivir. Taken together, most of the data so far support the administration of type I IFN early during infection to curb viral infection and lessen disease severity. Next to involvement of various cellular pathways, both on transcriptomic as well as proteomic level, we identified novel signatures in primary hAEC after infection with SARS-CoV-2. Strikingly, despite reduced viral replication in the presence of highly antiviral IFNα subtypes, infection with SARS-CoV-2 resulted in a downregulation of O-glycan processing. Mucus plays a vital role in protecting the respiratory tract from various factors, and serves as first line of defence against invading pathogens. Goblet cells secrete soluble mucus which major components are heavily O-glycosylated mucin glycoproteins^54^. Inflammatory conditions result in an increase of soluble and transmembrane mucins, and alteration of their glycosylation to boost mucosal defence^55,56^. Therefore, it is striking that we observed a consistent downregulation of various mucins upon SARS-CoV-2 infection. Some recent studies have highlighted the highest level of expression of ACE2 and TMPRSS2, entry factors utilized by SARS-CoV-2, in the nasal goblet and ciliated cells in healthy individuals, cells which are also associated with high MUC1 and MUC5A expression levels^57,58^. Therefore, it is likely that these cells represent the initial infection route for the virus. It is tempting to speculate that virus infection of these cells triggers mucin downregulation in order to impede cellular defence mechanisms. Interestingly, a significant proportion of COVID-19 patients represents with dry cough, indicating that downregulation of mucins could contribute to this clinical characteristic. In contrast, a recent study has described elevated MUC1 and MUC5AC protein levels in airway mucus of critical ill COVID-19 patients^59^. However, the authors speculated that elevated mucin levels could originate from detached and disrupted epithelial cells. It will be interesting to further analyze the role of mucins and their glycans during COVID-19 pathogenesis and study the influence of viral replication on mucin expression. In conclusion, in this study we provide a global characterization of the antiviral response of different IFNα subtypes on various levels and uncovered immune signatures which are able to significantly reduce SARS-CoV-2 infection as well as identify novel features after virus infection of primary cell types. Our study contributes to an enhanced understanding of the molecular landscape controlling SARS-CoV-2 infection and could thereby pave the way towards novel therapeutic approaches upon identification of key cellular pathways and factors involved in SARS-CoV-2 infection.

## Methods

### Stimulation with different human IFNα subtypes

IFNα subtypes were produced and purified as previously described ^7^. The activity of each subtype was determined using the human ISRE-Luc reporter cell line, a retinal pigment epithelial cell line transfected with a plasmid containing the Firefly Luciferase gene, stably integrated under control of the IFN-stimulation-response element (ISRE). Following stimulation with IFNα, chemiluminescence can be detected and used to calculate the respective activity in units against commercially available IFNα (PBL assays sciences, Piscataway, USA)^7^.

### End-point dilution assay

VeroE6 cells were seeded at a density of 10,000 cells per well in a 96-well plate and maintained in 200 μl DMEM (Gibco) supplemented with 10% fetal bovine serum (Sigma-Aldrich), L-glutamine (Gibco), penicillin and streptomycin (Gibco) overnight. The next day, 22 μl of virus stock or apical washes of hAEC were added to the first row of the plate (6 replicates). Then, the virus was diluted 1:10 by mixing the media and pipetting 22 μl to the next row repeatedly, followed by 72 h incubation in 37°C in a 5% CO_2_ atmosphere. Thereafter, the supernatant was aspirated and the cells were incubated in 100 μl of crystal violet solution (0.1 % crystal violet (Roth) in PBS, 10% ethanol, 0.37% formalin) for 5 min. Subsequently, the crystal violet solution was aspirated, cells were washed with PBS and the number of wells with intact or damaged cell layer were determined. The TCID_50_/mL was calculated by the Spearman & Kärber algorithm.

### IFN titration assay

VeroE6 cells were seeded at a density of 10,000 cells per well in a 96-well plate and maintained in DMEM supplemented with 10% fetal bovine serum, L-glutamine, penicillin and streptomycin overnight. Then, the medium was aspirated and serially diluted IFNα and IFNλ3 (R&D Systems) and virus with a final concentration of 350 PFU/mL were added to the cells in a total volume of 100 μl of cell culture media, followed by 72 h incubation in 37°C in a 5% CO_2_ atmosphere. Thereafter, the supernatant was aspirated and the cells were incubated in 100 μl of crystal violet solution (0.1 % crystal violet in PBS, 10% ethanol, 0.37% formalin) for 5 min. Subsequently, the crystal violet solution was aspirated, cells were washed with PBS and the number of wells with intact or damaged cell layer were determined.

The inhibitory concentration 50 (IC50) was calculated using GraphPad Prism 6.

### In-cell ELISA

The in-cell (ic) ELISA was performed based on the previously published protocol (Scholer et al., 2020). VeroE6 cells were seeded at a density of 20,000 cells per well in a 96-well plate and maintained in DMEM supplemented with 10% fetal bovine serum, L-glutamine, penicillin and streptomycin. At indicated time points, the medium was aspirated and serially diluted IFNα or the indicated concentrations of remdesivir and virus with a final concentration of 350 PFU/mL were added to the cells in a total volume of 100 μl, followed by 24 h incubation in 37°C in a 5% CO_2_ atmosphere. Thereafter, 100 μl of 8% ROTI®Histofix (Roth) (equals 4% of total PFA) were added for a minimum of 2 h at room temperature to fix the cells and inactivate the virus. Afterwards, the plate was washed thrice with PBS. The PBS was aspirated and 200 μl of freshly prepared permeabilization buffer (PBS, 1% Triton X-100 (Roth)) were added to the cells and the plate was incubated for 30 min at room temperature with constant shaking. Subsequently, the permeabilization buffer was aspirated and 200 μl of blocking buffer (PBS, 3% FBS) were added for 1 h. Then, the blocking buffer was aspirated and 50 μl of primary antibody solution (anti-SARS-CoV-2-NP (RRID: AB_2890255) 1:5000 diluted in PBS + 1% FBS) was added to each well. The plate was incubated overnight at 4°C. The next day, the primary antibody solution was aspirated and the plate was washed thrice with wash buffer (PBS, 0.05% Tween 20 (Roth)). Thereafter, 50 μl of the secondary antibody solution (Peroxidase-AffiniPure Goat Anti-Mouse IgG (H+L) (RRID: AB_10015289) 1:2000 in PBS, 1% FBS) was added to the wells and the plate was incubated for 2 h at room temperature. After the incubation period, the wells were washed 4 times with 250 μl wash buffer. Afterwards 100 μl of TMB substrate solution (BioLegend) were added and the plate was incubated about 20 min at room temperature in the dark. The reaction was stopped by addition of 100 μl 2N H_2_SO_4_ (Roth). The absorbance was measured at 450 nm with a reference wavelength of 620 nm using Spark® 10M multimode microplate reader (Tecan).

### Cell viability assay

To exclude cytotoxic effects of the compounds used in our assays, a cell viability assay was performed using the Orangu™ Cell Counting Solution (CELL guidance systems) according to the manufacturer’s instructions. The cells were seeded and treated equally to the protocol that was used before without any viral infection. Afterwards, 10 μl of Orangu™ Cell Counting Solution were added to each well and the plate was incubated for 2 h. Then, the absorbance was measured at 450 nm with Spark® 10M multimode microplate reader.

### Immunofluorescence

VeroE6 cells were seeded and treated as described for the in-cell ELISA. After incubation with the primary antibody solution, 50 μl secondary antibody solution (Goat IgG anti-Mouse IgG (H+L)-Alexa Fluor 488, MinX none 1:2000 (RRID: AB_2338840), Phalloidin CF647 1:100 (Biotium) in PBS + 1% FBS) were added to each well and the plate was incubated for 2 h at room temperature. Thereafter, the secondary antibody solution was aspirated and the cells were counterstained for 20 min at room temperature with 50 μl of DAPI solution (0.1 μg/mL DAPI (Sigma-Aldrich) in PBS). Subsequently, the plate was washed thrice with PBS and microscopically analyzed using Leica THUNDER Imager 3D Cell Culture.

### Infection of Human airway epithelial cells

Human airway epithelial cells (hAEC) were obtained from lung transplant donors post mortem (ethics of University Duisburg-Essen (18-8024-BO and 19-8717-BO)) or from explanted lungs (Ethics of Hannover medical school 3346/2016. Selection criteria for donors are listed in the Eurotransplant guidelines. hAECs from explanted lungs were cultured and differentiated as previously described ^60^ hAEC from lung transplant donors post mortem were obtained by the following protocol: During the adaptation of the donor lung, a small tracheal ring was removed and stored in PBS supplemented with antibiotics (penicillin 100 U/mL, streptomycin 100 μg/mL, 10 μg/mL ciprofloxacin (Kabi)). HAEC were isolated from the mucosa within 24 h after transplantation by enzymatic digestion (Protease XIV (Sigma Aldrich)) and scraping. Cells were expanded for 7-14 days in KSFM (keratinocyte-SF-medium (Gibco), supplemented with human epidermal growth factor (Gibco) (2.5 ng/mL), bovine pituitary extract (Gibco) (BPE 25 μg/mL, Gibco), isoproterenol (Sigma-Aldrich) (1μM), Penicillin, Streptomycin, Ciprofloxacin, Amphotericin B (PanBiotech) (2,5 μg/mL)) and after trypsinization stored in liquid nitrogen (10% DMSO, 90% KSFM+BPE 0,3mg/mL). All plastic surfaces during hAEC isolation and air liquid interface (ALI) culture were coated with human fibronectin (PromoCell) (5 μg/mL), type I bovine collagen (Advanced BioMatrix) (PureCol 30 μg/mL) and BSA (10 μg/mL). For ALI cultures, cells were thawed, expanded in KSFM for 5-7 days and transferred to transwell inserts (PE Membrane, 12 well plates, 0.4 μm pore size, Corning). A monolayer hAECs were grown submerged in S/D Media (1:1 mixture of DMEM (StemCell) and BEpiCM-b (ScienCell), supplemented with Penicillin and Streptomycin, HEPES (Gibco) (12.5mL/l, 1M), 1x Bronchial Epithelial Cell Growth Supplement (ScienCell), and EC-23 (Tocris) (5mM) until they reached confluency. Apical media was removed and cell differentiation was induced under air exposure for 2 weeks. Infection was started after cells were fully differentiated measured by movement of cilia, secretion of mucus and transepithelial electrical resistance (>1000Ω/cm2).

Fully differentiated hAECs were washed with HBSS apically for 10 min before infection. For SARS experiments, the cells were infected apically with 30,000 PFU diluted in HBSS, for Influenza, the cells were apically infected with Influenza A virus H1H1 strain A/Puerto Rico/34 (PR8) at 0.1 MOI in 200 μl HBSS. The cells were incubated with the inoculum for 1 h in 33°C in a 5 % CO2 atmosphere. Thereafter, the inoculum was aspirated and the cells were washed thrice with 150 μl of HBSS for 10 min. The last wash was collected and stored at −80 °C as 0 h sample. At the indicated time points, cells were washed apically for 10 min and the washes were subjected to an end-point dilution assay or to a plaque titration assay as described for SARS-CoV-2 and Influenza, respectively.

Treatment of hAECs was performed by adding the indicated amounts of IFNs or remdesivir directly to the cell culture medium on the basolateral side of the cells.

For the isolation of RNA, cells were lysed using Qiagen RLT buffer (Qiagen) supplemented with 1% β-mercaptoethanol (Sigma-Aldrich).

### Viral mRNA quantification

Total RNA was purified from hAECs and VeroE6 cells using the RNeasy Mini Kit (Qiagen) according to manufacturer’s instructions with preceding DNase I digestion with the RNase-Free DNase Set (Qiagen).

To determine relative SARS-CoV-2 M- or N-gene expression, 500 ng of total RNA were reverse transcribed using the PrimeScript™ RT Master Mix (Takara). Promega’s GoTaq® Probe qPCR Master Mix was used according to the manufacturer’s instructions with gene specific primers and probes (see Extended data table 7). RT-qPCR was performed on a LightCycler® 480 II (Roche) instrument, with the following conditions: initial denaturation was 2 min at 95 °C and a ramp rate of 4.4 °C/s, followed by 40 cycles of denaturation for 15 seconds at 95 °C and a ramp rate of 4.4 °C/s and amplification for 60 seconds at 60°C and a ramp rate of 2.2 °C/s. To assess M- and N-gene copy numbers, the M- and N-gene were partially cloned into pCR™2.1 (ThermoFisher Scientific) or pMiniT 2.0 (NEB), respectively, and a 1:10 plasmid dilution series was used as a reference.

### IAV plaque assay

MDCK-II cells were seeded in 6 well plates, and cultured in DMEM supplemented with 5% FBS and 1% Penicillin-Streptomycin until 100% confluent. On the day of infection, 10-fold dilutions of apical washes were prepared in infection-PBS (PBS supplemented with 1% Penicillin-Streptomycin, 0.01% CaCl2, 0.01% MgCl2 and 0.2% BSA). Cells were washed once with infection-PBS, infected with 500 μl of diluted samples (virus inoculum), and were incubated at 37°C, 5% CO2 for 30 min. The inoculum was removed, and the infected monolayer was overlaid with plaque medium (prepared immediately before use by mixing 14.2% 10X MEM (Gibco), 0.3% NaHCO3, 0.014% DEAE-Dextran (Sigma-Aldrich), 1.4% 100X Penicillin-Streptomycin, 0.3% BSA, 0.9% Agar, 0.01% MgCl2, 0.01% CaCl2, 0.15 mg TPCK-Trypsin (Sigma). Plates were kept at room temperature until the agar solidified, and were incubated upside down at 37°C, 5% CO2 for 72h. Plaques were quantified in terms of infectious IAV particles, and were represented as PFU/mL.

### ISG expression

500,000 VeroE6 cells were seeded and stimulated with 1000 U/mL of IFNα subtypes 5, 7, 16, or 1000 ng/mL IFNλ3 for 16 h. Afterwards, the cells were lysed using DNA/RNA Shield for RNA isolation.

RNA was isolated from cell lysates with Quick-RNA™ Miniprep Kit (Zymo Research) according to the manufacturer’s instruction.

CDNA was synthesized from isolated RNA using cDNA Synthesis Super Mix (Bimake) according to the manufacturer’s instructions. ISG expression levels were quantified by qPCR with Luna® Universal qPCR Master Mix and the respective primer pairs (see Extended data table 6). Expression levels were normalized by 2-ΔΔCT method^61^ using GAPDH as reference gene.

### Proteomics sample preparation

Cells were washed with ice cold PBS and harvested in urea buffer (30 mM Tris HCl, 7 M Urea, 2 M Thiourea, 0.1% NaDOC, pH 8.5). Cells were centrifuged for 15 min at 16,100 x g and 4 °C and the supernatant was further processed.

Tryptic digestion was performed on 20 μl cell lysate. Disulfide bonds were reduced by adding final 5 mM DTT (Dithiothreitol) for 15 minutes at 50 °C before thiols were alkylated by final 15 mM IAA (iodoacetamide) for 15 min in the dark. Hydrophilic and hydrophobic Cytiva Sera-Mag Carboxyl-Magnet-Beads (GE Healthcare) were mixed 1:1 and 2 μl beads (25 μg/μl) were added per samples. The samples were filled up to 70% ACN (acetonitrile) and incubated for 15 min to ensure protein binding to the beads. Subsequently, beads were washed two times with 70% EtOH followed by washing with 100% ACN. Beads were resuspended in 100 mM ammonium bicarbonate carbonate containing 0.2 μg trypsin (SERVA) per sample and incubated overnight at 37 °C. The peptides were transferred into a new reaction tube, vacuum dried and dissolved in 0.1 % TFA (trifluoroacetic acid).

### LC-MS/MS Analysis

400 ng tryptic peptides per sample were analyzed using an Ultimate 3000 RSLCnano HPLC (Dionex) coupled to a Q Exactive HF Orbitrap (Thermo Fisher Scientific). Samples were pre-concentrated on a C18 trap column (Acclaim PepMap 100; 100 μm × 2 cm, 5 μm, 100 Å; Thermo Fisher Scientific) within seven minutes at a flow rate of 30 μL/min with 0.1 % trifluoric acid and subsequently transferred to a Nano Viper C18 analytical column (Acclaim PepMap RSLC; 75 μm × 50 cm, 2 μm, 100 Å; Thermo Fisher Scientific). Peptide separation was performed by a gradient from 5% - 30% solvent B over 120 minutes at 400 nL/min (solvent A: 0.1% formic acid; solvent B: 0.1% formic acid, 84% acetonitrile). Full-scan mass spectra were acquired in profile mode at a resolution of 70,000 at 400 m/z within a mass range of 350 – 1400 m/z. The 10 highest abundant peptide ions were fragmented by HCD (NCE [normalized collision energy] = 27) and MS/MS spectra were acquired at a resolution of 35,000.

### Proteomics Data Analysis

Peptide identification and quantification were performed using MaxQuant (v.1.6.17) searching UniProtKB/SwissProt (2020_05, 563,552 entries) restricted to either Homo sapiens or Homo sapiens and SARS-CoV-2. Search parameters were default, LFQ was used for peak quantification and normalization was enabled. Peptides were considered for quantification irrespective of modifications. Match between runs was enabled when the analysis was performed considering human proteins only. Statistical data analysis was conducted using R (v.3.6.2). Differences between the experimental groups were assessed using t-tests (paired, two-sided) and proteins quantified in minimum 3 of 4 donors per group with minimum 2 unique peptides, a p-value ≤ 0.05 and a ratio of mean abundances ≥ 1.5 or ≤ 0.67 were considered statistically significant. Proteins that were quantified in one experimental group but not detected at all in an opposed group were defined as On-Offs between these groups. GO annotation and enrichment analyses were performed using STRING (v.11). Data visualization was done using R and Cytoscape (v.3.8.2).

### Data availability

The authors declare that the data supporting the findings of this study are available within the article and its Extended Data files or are available on request. The mass spectrometry proteomics data have been deposited at the ProteomeXchange consortium via the PRIDE partner repository with the dataset identifier PXD000XXX.

### Transcriptomics

Quality and integrity of total RNA was controlled on 5200 Fragment Analyzer System (Agilent Technologies)). The RNA sequencing library was generated from 50 ng total RNA using NEBNext® Single Cell/Low Input RNA Library to manufacture’s protocols. The libraries were treated with Illumina Free Adapter Blocking and were sequenced on Illumina NovaSeq 6000 using NovaSeq 6000 S1 Reagent Kit (100 cycles, paired end run 2× 50 bp) with an average of 3 ×10^7^ reads per RNA sample.

### Transcriptomic analysis

FASTQ files of RNA sequencing files were imported into the Array Studio software v10.2.5.9 (QIAGEN, Cary, NC, USA) package for further data analysis. All FASTQ files were aligned to the gene model Ensembl v96 and to the reference library Human B38 using the proprietary OmicSoft Aligner OSA^62^. Differential gene expression of each condition was assessed using DESeq2^63^. Differentially expressed genes were sent to Ingenuity Pathway Analysis (IPA) (https://digitalinsights.qiagen.com/products-overview/discovery-insights-portfolio/analysis-and-visualization/qiagen-ipa/) for biological analysis using the cutoffs: p-value <0.05, fold change (fc) >|1.5| and mean counts min>5. IPA statistics is based on two outputs. A p-value derived from a right-tailed Fisher’s Exact Test estimates the probability that the association between a function or pathway and a set of molecules might be due to random chance but does not consider directional changes. This is, however, predicted for a disease and/or function, canonical pathway, or upstream regulator (activation or inhibition) by the activation z-score algorithm. The z-score describes the number of standard deviations data lies above or below the mean. A z-score >2 was considered significantly increased whereas a z-score<−2 was considered significantly decreased^64^. We performed an expression analysis to evaluate transcriptomic changes for Canonical Pathways in each of the comparison IFN vs mock^64^.

### Statistical analysis

Differences in transformed data were tested for significance using GraphPad Prism v8.4.2 for Windows (GraphPad). Statistically significant differences between the IFNα-treated groups and the untreated group were analyzed using Ordinary One-Way ANOVA analysis with Dunnetts’s multiple comparison test. P values < 0.05 were considered significant.

## Acknowledgements

We would like to thank all member of the Department for Molecular and Medical Virology of the Ruhr University Bochum for helpful suggestions and discussion. We thank Kristin Fuchs and Birgit Zülch for excellent technical assistance. We want to thank the Westdeutsche Biobank Essen (WBE, University Hospital Essen, University of Duisburg-Essen, Essen, Germany; approval WBE-ref. 20-WBE-102) for providing Human biological samples.

## Author Contributions

S.P., K.S., E.S., U.D., D.T., designed the project. J. S., T.L.M., K.S., C.E., N.H., Z.K., S.H., S.K., L.B., B.W., H.B., A.K. performed and analysed experiments. D.T. T.B., K.Sch., J.N.B., and M.E. performed statistical analysis and data analysis. T.P., B.S., J.C., Z.Y., C.T., V.T.K.L.T., M.T. and S.L. contributed to the design and implementation of the research. S.P., and K.S., wrote the manuscript, and all authors contributed to editing.

## Competing interest declaration

The authors declare no competing interests. J.N.B. is an employee of QIAGEN, Inc. (no conflict of interest).

## Funding

K.S. was supported by a grant from the DFG (SU1030-2-1) and Stiftung Universitätsmedizin Essen.

S.P. was supported by a grant from the Federal Ministry of Education and Research (BMBF) under grant agreement no. 01KI2058.

K.Sch. was supported by the German Network for Bioinformatics Infrastructure (de.NBI), a project of the German Federal Ministry of Education and Research (BMBF) [FKZ 031 A 534A]. M.E. works within the research building Center for Protein Diagnostics (PRODI), funded by North Rhine-Westphalia state and German Federal funds.

L.B. and S.L. received funding from the BMBF (grant no. 01KI20218, CoIMMUNE), the Netzwerk Universitäts Medizin COVID-19 (NUM-COVID-19) within the Network Organo-Strat (grant no.01KX2021), the fund “Innovative Medical Research of the University Muenster Medical School” (grant no. BR111905; to L.B.) and from the German Research Foundation (DFG) for TP6 as part of the CRU 342 (to L.B. and S.L.) and DFG Grants SFB1009 B13 (to SL).

## Extended Data Figures and legends

**Extended Data Fig. 1:**
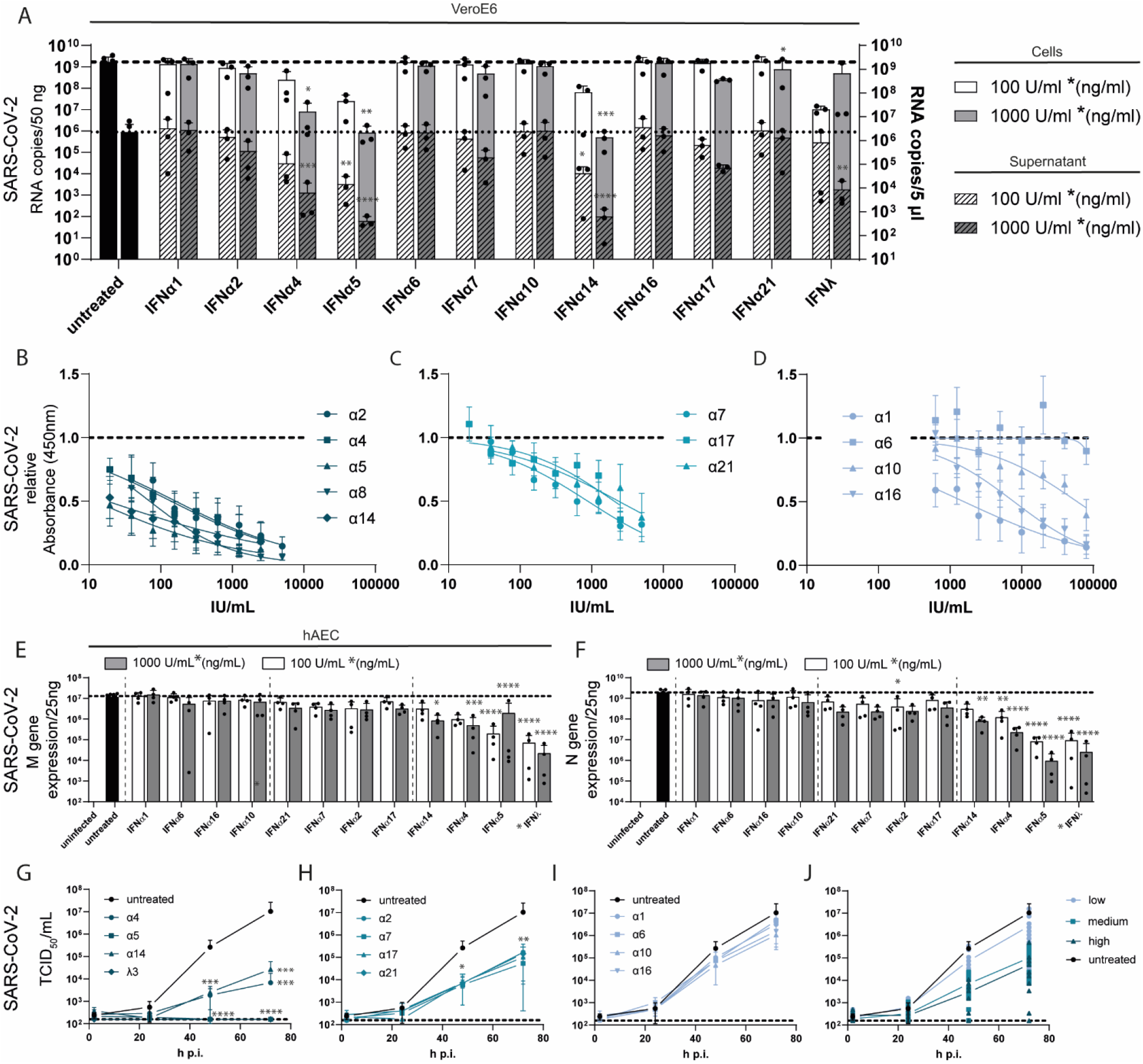
Treatment with IFNα subtypes reveals distinct antiviral effects against SARS-CoV-2. (A) Antiviral activity of IFNα subtypes and IFNλ3 against SARS-CoV-2 were analysed in VeroE6 cells and cell supernatant by qRT-PCR. IFNα subtypes were titrated against SARS-CoV-2 on VeroE6 cells by ic-ELISA assay and the IFNs were grouped as mean values in high (B), medium (C) and low (D) antiviral pattern. Antiviral activity of IFNα subtypes and IFNλ3 in SARS-CoV-2-infected primary hAECs at 72 h p.i. determined by qRT-PCR analysis of *M gene* (E) and *N gene* (F). Kinetics of the antiviral activity of IFNs by TCID_50_ assay grouped into high (G), medium (H) and low (I) antiviral pattern and the mean values of each group are plotted in (J). Mean values of high (I) and low/not (J) antiviral IFNs are shown. Mean values ± SEM are shown. (A-D) n=3 (E-J) n=4. A: RNA copies /50ng: * p=0.0228 (1000U/mL IFNα4); * p=0.0110 (1000U/mL IFNα21); ** p=0.0021 (1000U/mL IFNα5); *** p=0.0008 (1000U/mL IFNα14); RNA copies/5μl : * p=0.0106 (100U/mL IFNα14); ** p=0.0050 (100U/mL IFNα5); ** p=0.0017 (1000ng/mL IFNλ3); *** p=0.0002 (1000U/mL IFNα4); **** p<0.0001 (1000U/mL IFNα5, α14) E: 100 U/mL (ng/mL): **** p<0.0001 (IFNα5, λ3); 1000 U/mL (ng/mL): * p=0.0184 (IFNα14); *** p=0.0003 (IFNα4) **** p<0.0001 (IFNα5, λ3); F: 100 U/mL (ng/mL): * p=0.0289 (IFNα2); ** p=0.0032 (IFNα4)**** p<0.0001 (IFNα5, λ3); 1000 U/mL (ng/mL): ** p=0.0019 (IFNα14); **** p<0.0001 (IFNα4, α5, λ3); G: 48h: * p= 0.0120 (IFNα2); *** p= 0.0001 (IFNα4); *** p= 0.0002 (IFNα14); **** p<0.0001 (IFNα5, λ3); 72h: ** p= 0.0034 (IFNα2); **** p<0.0001 (IFNα4, α5, α14, λ3) H: 48h: * p= 0.0278 (IFNα7); * p= 0.0179 (IFNα21);72h: ** p= 0.0011 (IFNα7); ** p= 0.0031 (IFNα17); ** p= 0.0037 (IFNα21)

**Extended Data. Fig. 2:**
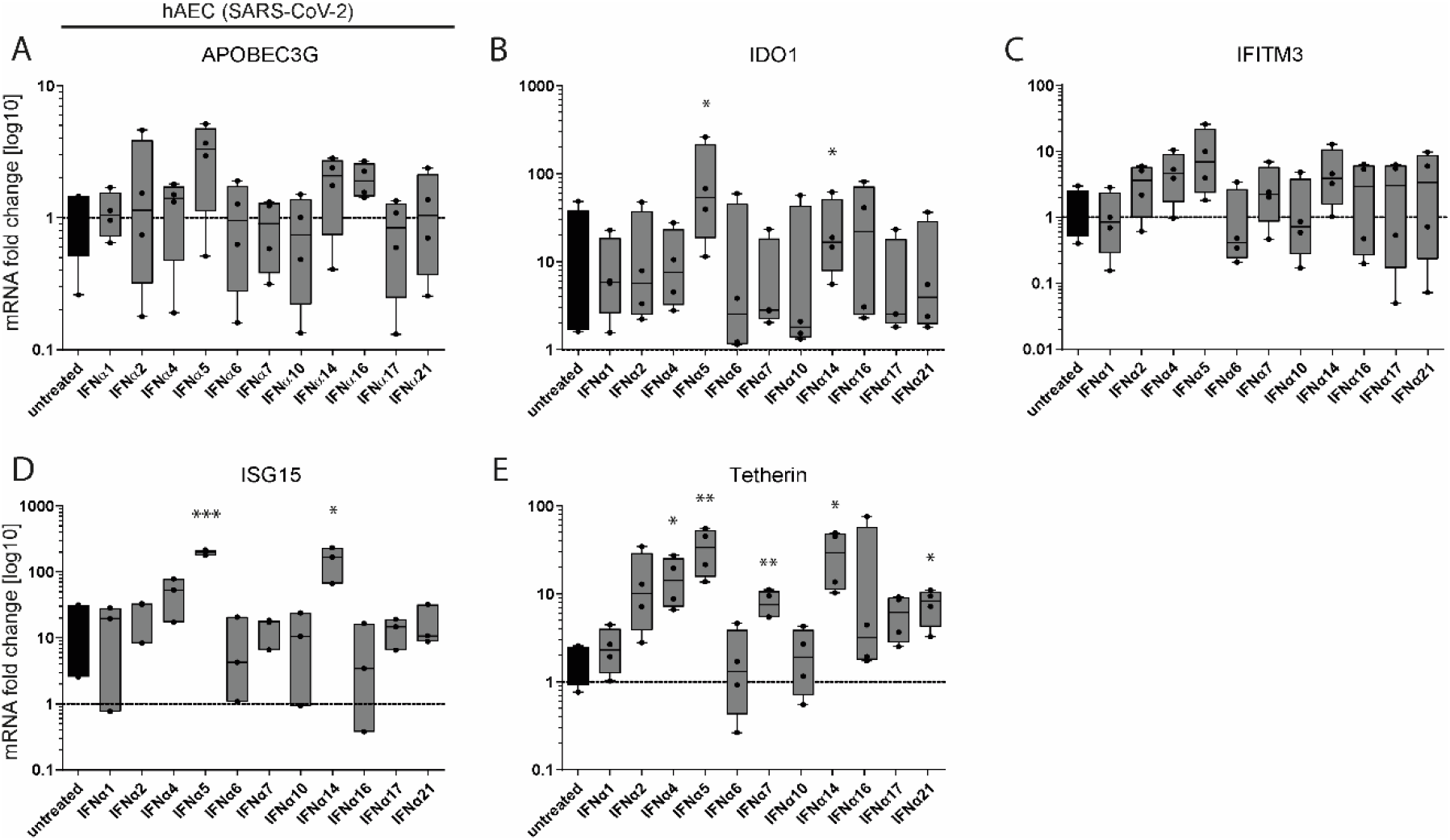
ISG induction upon IFNα subtype stimulation. (A-E) mRNA expression of different ISGs in IFN-treated SARS-CoV-2-infected primary hAECs at 72 h p.i. determined by qRT-PCR analysis. Mean values of mRNA expression is shown as fold change compared to untreated control. (A-E) n=4. B: * p=0.0392 (IFNα5); * p=0.0460 (IFNα14); I: * p=0.0198 (IFNα14); *** p=0.0004 (IFNα5); J: * p=0.0200 (IFNα4); * p=0.0197 (IFNα14); * p=0.0247 (IFNα21); ** p=0.0087 (IFNα5); ** p=0.0079 (IFNα7)

**Extended Data Fig. 3:**
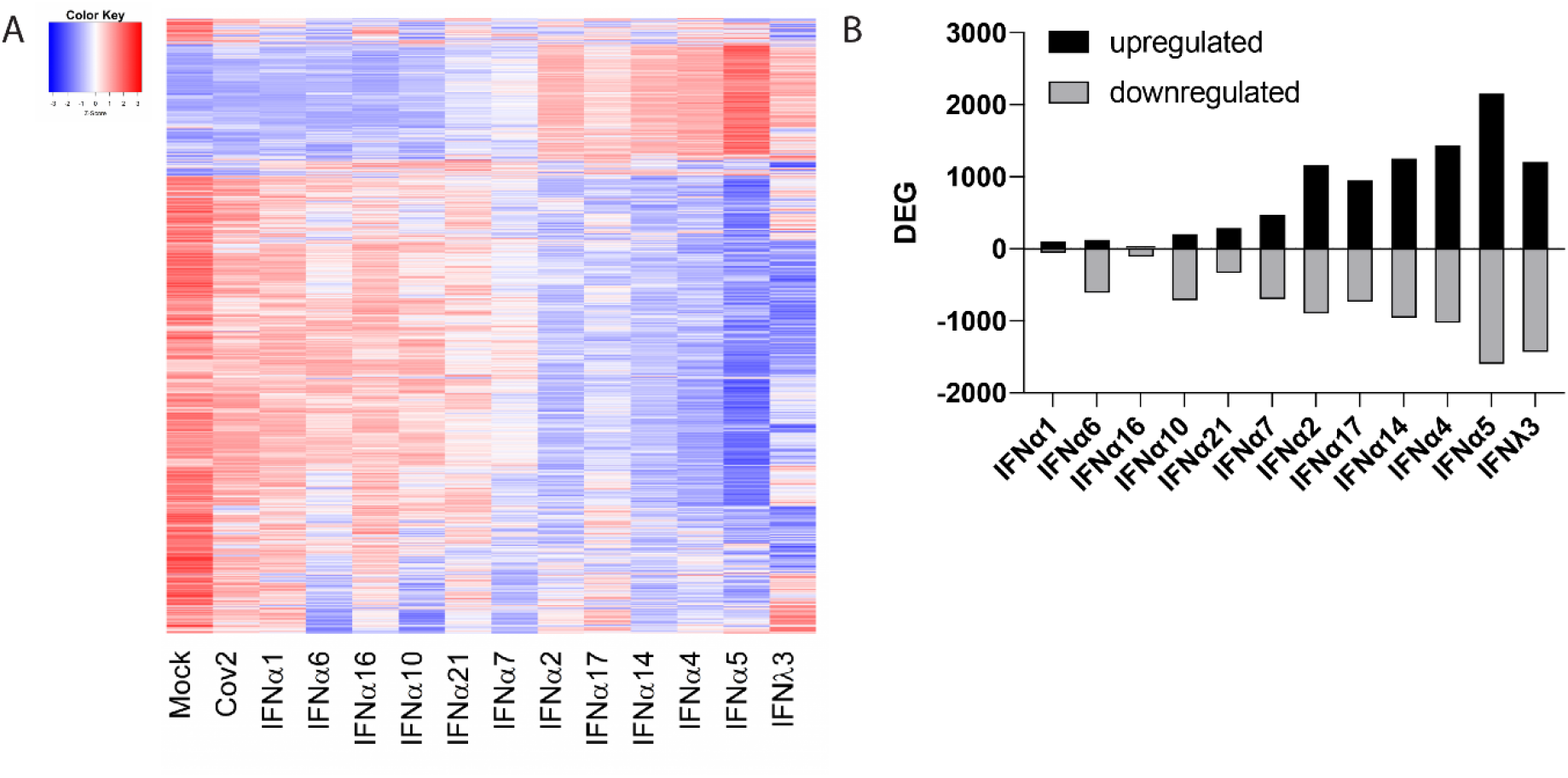
Transcriptomic analysis display IFNα subtype specific gene signatures. (A) Numbers of up- and downregulated DEGs of IFN-treated compared to untreated hAECs (4 donors) shown as bars. (B)Transcriptomic analyses of IFN-treated (16 hours post treatment) or SARS-CoV-2-infected (18 hours post infection) hAECs. Heat maps displaying differentially expressed genes (DEG) from at least one comparison of an IFN vs. Mock.

**Extended Data Fig. 4:**
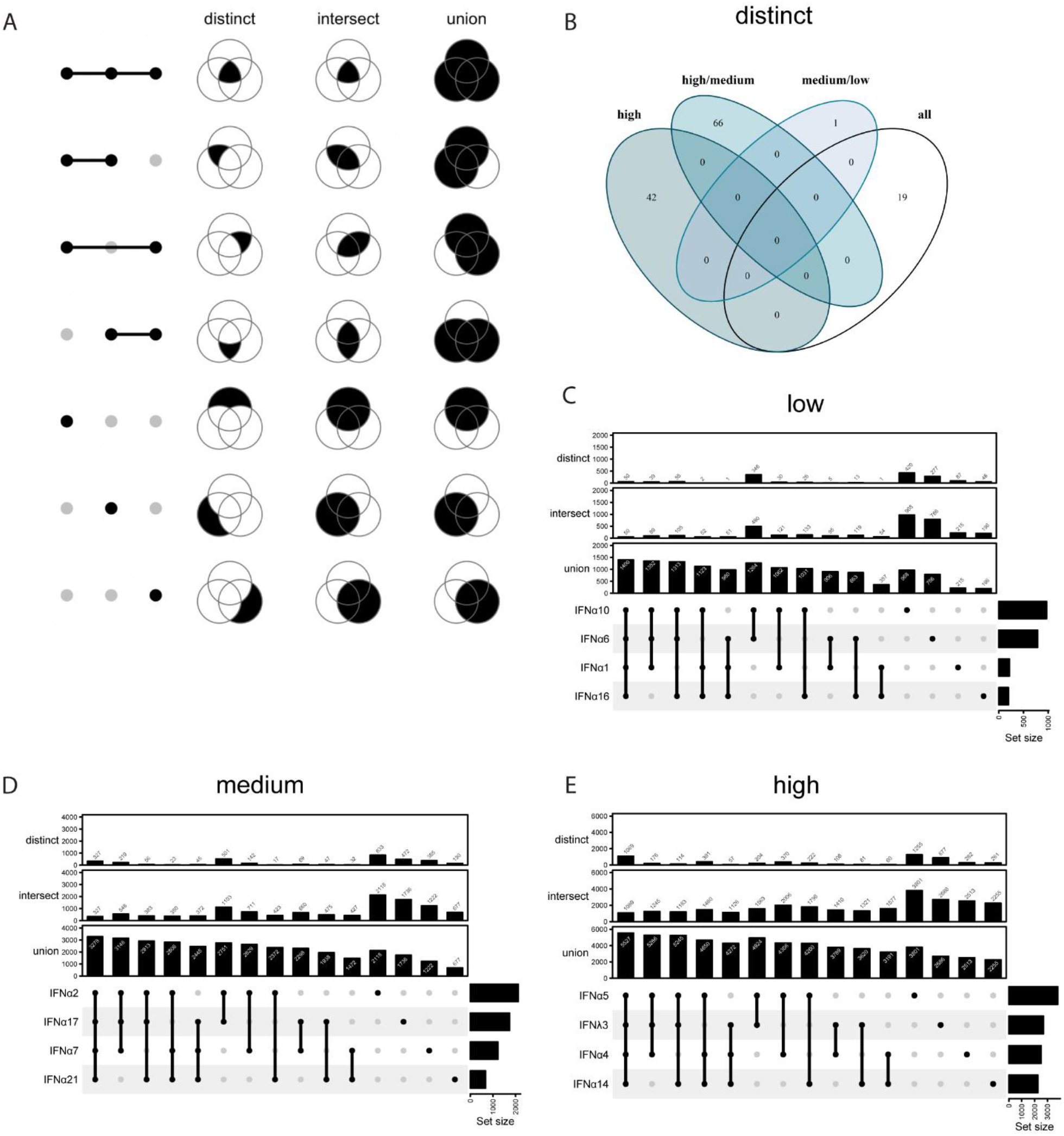
Transcriptomic analysis reveal different patterns of distinct, intersect and union genes. (A-E) UpSet plots to summarize distinct, intersect and union differentially expressed genes (DEG) of IFN-treated (16 hours post treatment) hAECs (4 donors). (A) Schematic depiction of distinct, intersect and union DEGs. (B) Venn diagram of distinct DEGs expressed by all high, medium and low antiviral IFNs. (C) UpSet plots showing distinct, intersect and union DEGs of low (C), medium (D) and high (E) antiviral IFNs. Numbers of individually or group-specific DEGs are shown as bars and numbers. The bottom right horizontal bar graph labelled Set Size shows the total number of DEGs per treatment.

**Extended Data Fig. 5:**
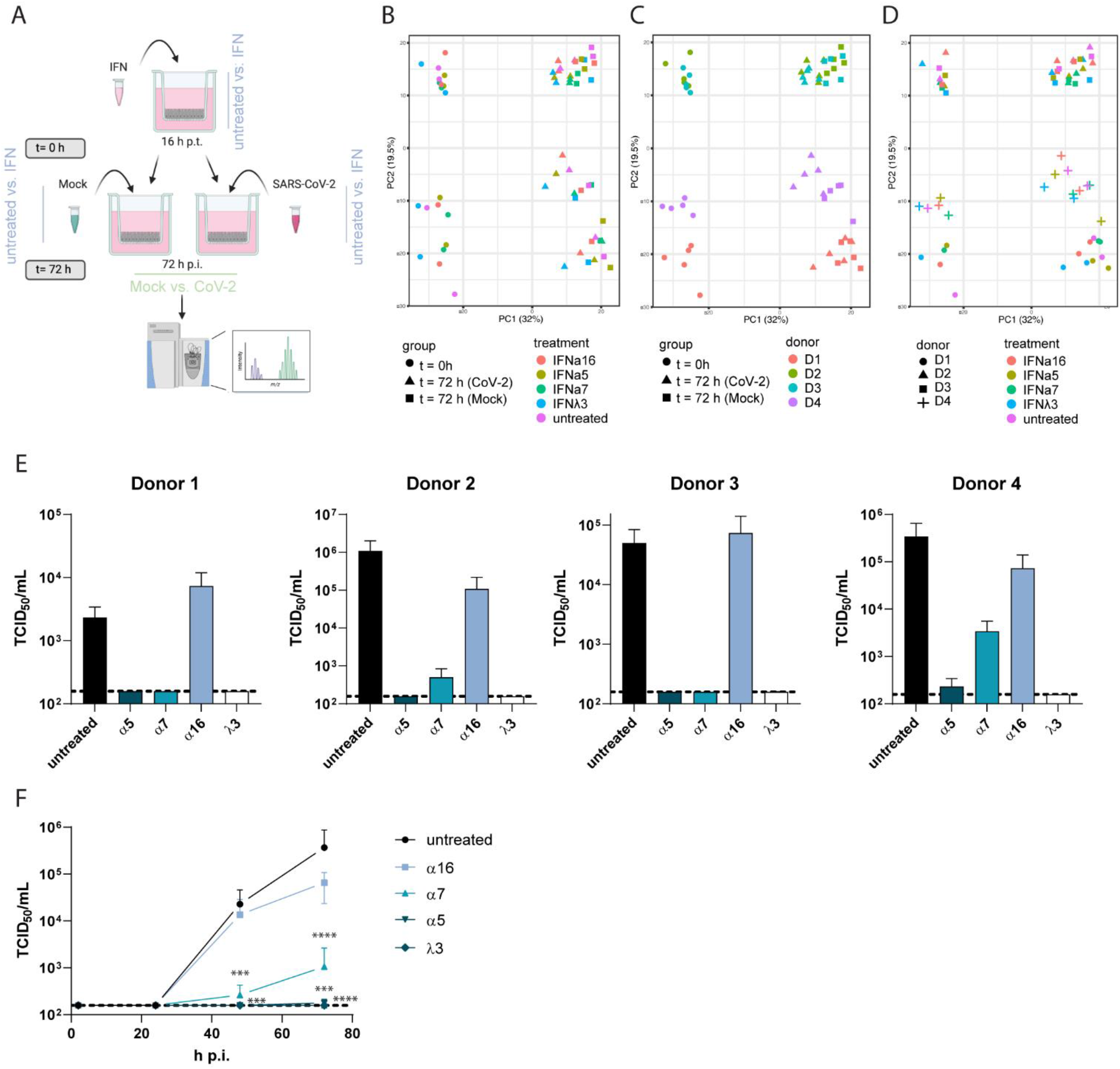
Proteomic analysis highlights key cellular mediators. (A-D) Proteomic analysis of IFN-treated and/or SARS-CoV-2-infected hAECs. (A) Schematic depiction (B-D) Principal component analysis (PCA) of hAEC proteomics. (B) The first two principal components (PCs) are plotted and shaped/coloured according to group and IFN-treatment (B); to group and individual donors (C) or to individual donors and IFN-treatment (D). PCA was performed using all proteins without missing values. Percentage of variation accounted for by each principal component is shown in brackets with the axis label. (E) Antiviral activity of IFNα subtypes and IFNλ3 in SARS-CoV-2-infected primary hAECs of 4 individual donors used for proteomic analysis at 72 h p.i. determined by TCID_50_ assay. (F) Kinetics of the antiviral activity of selected IFNs by TCID_50_ assay in SARS-CoV-2-infected primary hAECs of 4 individual donors used for proteomic analysis shown as mean values + SEM. D: 48h: *** p= 0.0003 (IFNα7); *** p= 0.0001 (IFNλ3, IFNα5); 72h: *** p= 0.0001 (IFNα5); *** p= 0.0006 (IFNα7); **** p<0.0001 (IFNλ3)

**Extended Data Fig. 6:**
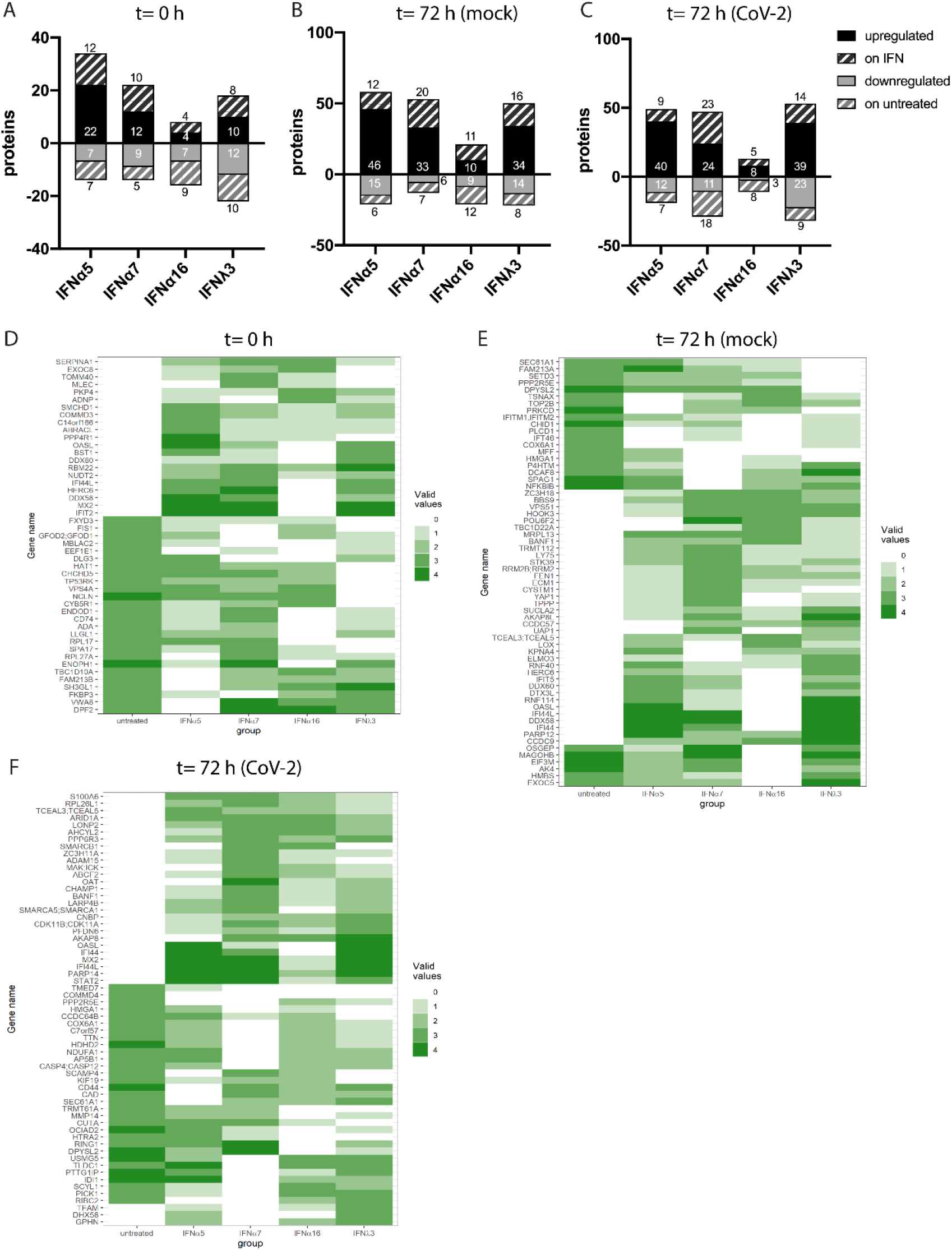
Proteomic analysis results in differential switched on/off proteins Proteomic analysis of IFN-treated and/or SARS-CoV-2-infected hAECs. (A) Differentially regulated or induced (on IFN compared to untreated) proteins in IFN-stimulated hAECs at t=0h (A), at t=72h (mock) (B) or at t=72h (CoV-2) (C). Heatmaps of on-off regulated proteins in IFN-stimulated hAECs at t=0 h (D), at t=72 h (mock) (E) or at t=72 h (CoV-2) (F).

**Extended Data Fig. 7:**
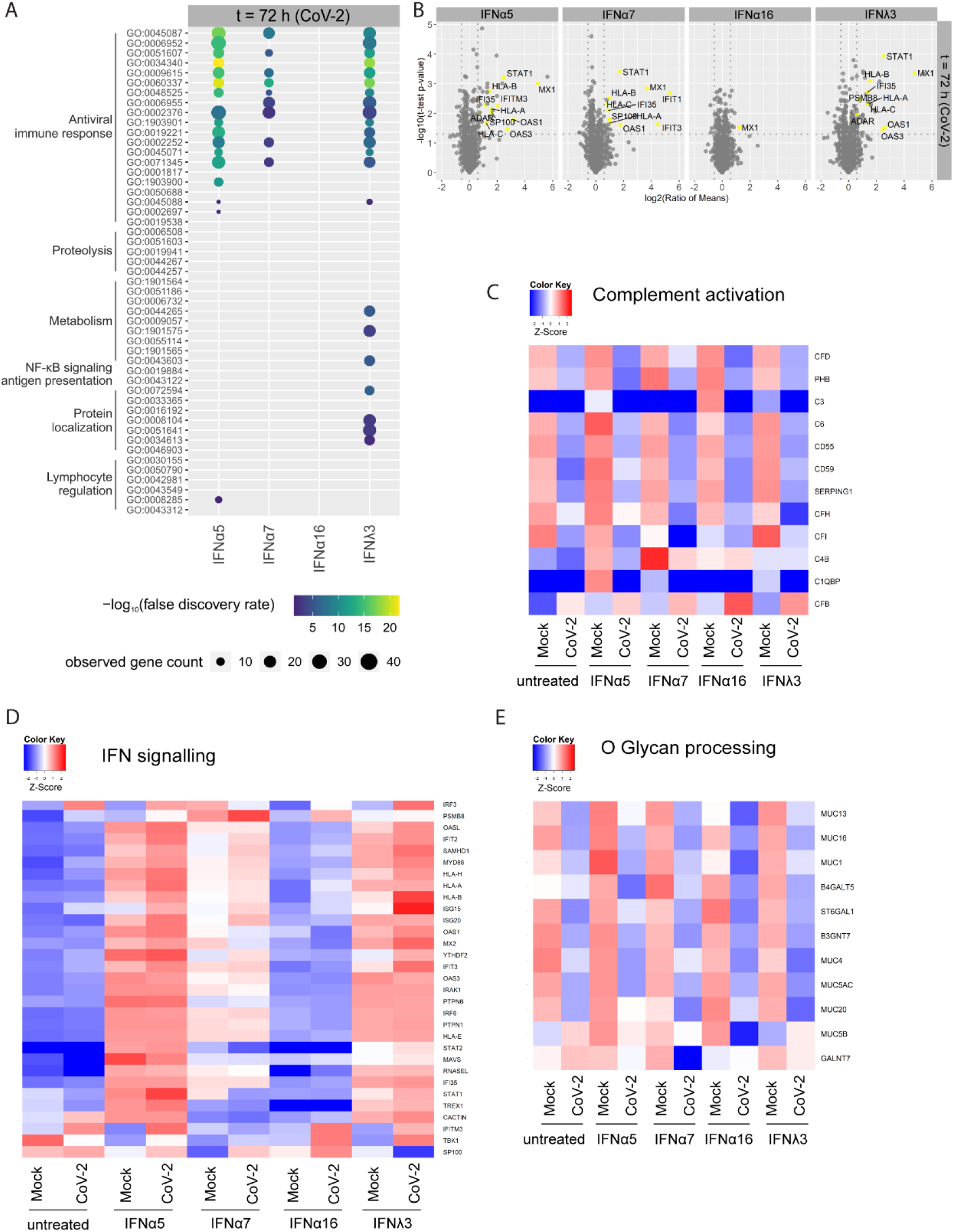
IFN signature did not change upon SARS-CoV-2 infection. (A-E)Proteomic analysis of IFN-treated and/or SARS-CoV-2-infected hAECs. (A) Biological processes induced by IFNs in SARS-CoV-2-infected hAECs at 88 h p. treatment (t=72 h (CoV-2)). (B) Volcano plots of IFN-treated SARS-CoV-2-infected hAECs (t=72 h (CoV-2)) Detected ISGs are coloured yellow. Heatmaps displaying differentially expressed proteins which are associated with complement activation (C) IFN signalling (D) and O glycan processing (E). Comparisons of IFN-treated mock or SARS-CoV-2 infected hAECs at 72 h p.i. are depicted. n=4.

**Extended Data Fig. 8:**
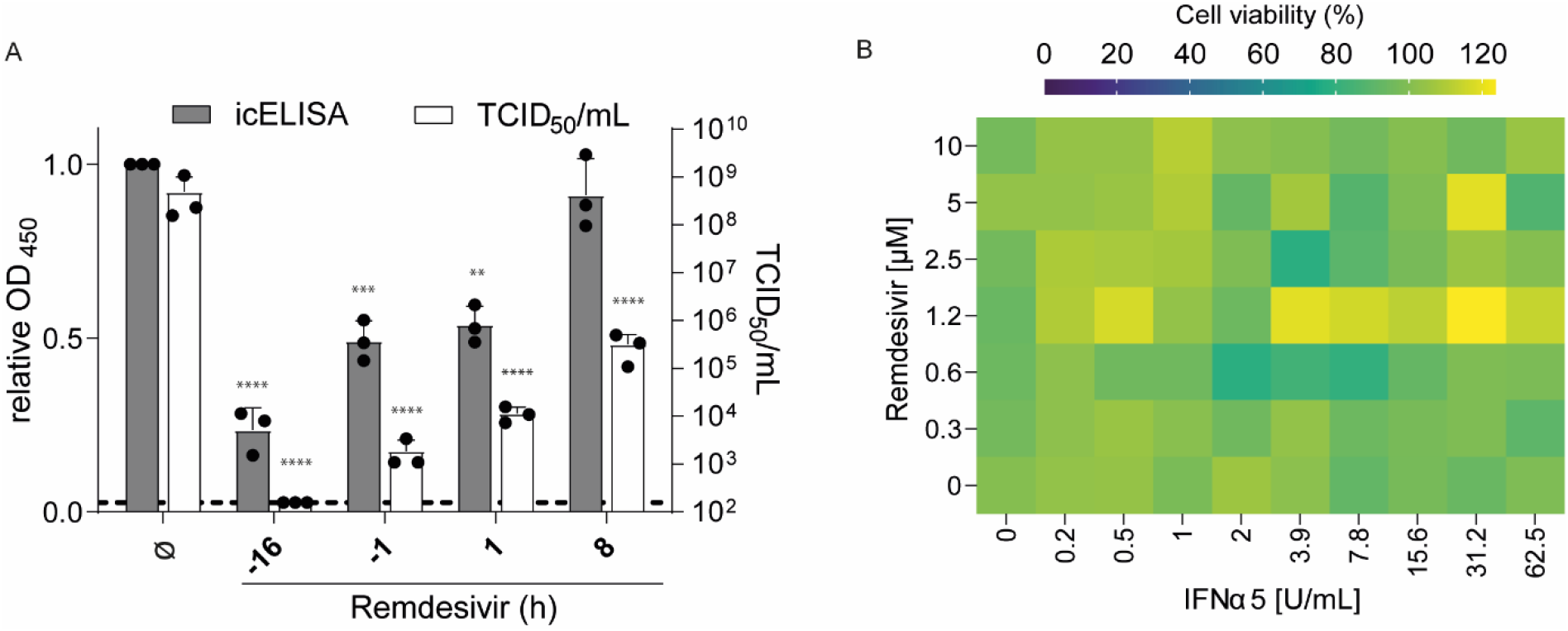
Therapeutic potential of combination treatment. (A) Pre- and post-treatments of VeroE6 cells with remdesivir analysed by icELISA (black bars) and TCID50 assay (white bars) shown as mean values + SEM. (B) Single and combined pre-treatments of IFNα5 and remdesivir in SARS-CoV-2 infected VeroE6 cells. Cell viability (%) normalised to untreated control (100%) is shown as heatmap. n=3. B: icELISA (grey bars) ** p=0.0024 (+1); *** p=0.0009 (−1); **** p<0.0001 (−16); TCID50/mL (white bars) **** p<0.0001 (−16, −1, +1, +8)

## Notes

### Competing Interest Statement

The authors have declared no competing interest.

## References

1 Hardy, M. P., Owczarek, C. M., Jermiin, L. S., Ejdeback, M. & Hertzog, P. J. Characterization of the type I interferon locus and identification of novel genes. Genomics 84, 331–345, doi:10.1016/j.ygeno.2004.03.003 (2004).

2 Mesev, E. V., LeDesma, R. A. & Ploss, A. Decoding type I and III interferon signalling during viral infection. Nat Microbiol 4, 914–924, doi:10.1038/s41564-019-0421-x (2019).

3 Platanias, L. C. Mechanisms of type-I- and type-II-interferon-mediated signalling. Nat Rev Immunol 5, 375–386 (2005).

4 Wittling, M. C., Cahalan, S. R., Levenson, E. A. & Rabin, R. L. Shared and Unique Features of Human Interferon-Beta and Interferon-Alpha Subtypes. Front Immunol 11, 605673, doi:10.3389/fimmu.2020.605673 (2020).

5 Chen, J. et al. Functional Comparison of Interferon-alpha Subtypes Reveals Potent Hepatitis B Virus Suppression by a Concerted Action of Interferon-alpha and Interferon-gamma Signaling. Hepatology 73, 486–502, doi:10.1002/hep.31282 (2021).

6 Dickow, J. et al. Diverse Immunomodulatory Effects of Individual IFNalpha Subtypes on Virus-Specific CD8(+) T Cell Responses. Front Immunol 10, 2255, doi:10.3389/fimmu.2019.02255 (2019).

7 Lavender, K. J. et al. Interferon Alpha Subtype-Specific Suppression of HIV-1 Infection In Vivo. J Virol 90, 6001–6013, doi:10.1128/JVI.00451-16 (2016).

8 Harper, M. S. et al. Interferon-alpha Subtypes in an Ex Vivo Model of Acute HIV-1 Infection: Expression, Potency and Effector Mechanisms. PLoS pathogens 11, e1005254, doi:10.1371/journal.ppat.1005254 (2015).

9 Gibbert, K., Schlaak, J., Yang, D. & Dittmer, U. IFN-alpha subtypes: distinct biological activities in anti-viral therapy. British journal of pharmacology 168, 1048–1058, doi:10.1111/bph.12010 (2013).

10 Song, J. et al. Different antiviral effects of IFNalpha subtypes in a mouse model of HBV infection. Sci Rep 7, 334, doi:10.1038/s41598-017-00469-1 (2017).

11 Schoggins, J. W. A diverse range of gene products are effectors of the type I interferon antiviral response. Nature 472, 481–485 (2011).

12 Lavoie, T. B. et al. Binding and activity of all human alpha interferon subtypes. Cytokine 56, 282–289, doi:10.1016/j.cyto.2011.07.019 (2011).

13 Jaks, E., Gavutis, M., Uze, G., Martal, J. & Piehler, J. Differential receptor subunit affinities of type I interferons govern differential signal activation. J Mol Biol 366, 525–539 (2007).

14 Tomasello, E., Pollet, E., Vu Manh, T. P., Uze, G. & Dalod, M. Harnessing Mechanistic Knowledge on Beneficial Versus Deleterious IFN-I Effects to Design Innovative Immunotherapies Targeting Cytokine Activity to Specific Cell Types. Frontiers in immunology 5, 526, doi:10.3389/fimmu.2014.00526 (2014).

15 Perrillo, R. Benefits and risks of interferon therapy for hepatitis B. Hepatology 49, S103–111, doi:10.1002/hep.22956 (2009).

16 Tan, G., Song, H., Xu, F. & Cheng, G. When Hepatitis B Virus Meets Interferons. Front Microbiol 9, 1611, doi:10.3389/fmicb.2018.01611 (2018).

17 Chan, H. L. Y. et al. Peginterferon lambda for the treatment of HBeAg-positive chronic hepatitis B: A randomized phase 2b study (LIRA-B). J Hepatol 64, 1011–1019, doi:10.1016/j.jhep.2015.12.018 (2016).

18 Kotenko, S. V., Rivera, A., Parker, D. & Durbin, J. E. Type III IFNs: Beyond antiviral protection. Semin Immunol 43, 101303, doi:10.1016/j.smim.2019.101303 (2019).

19 Lazear, H. M., Schoggins, J. W. & Diamond, M. S. Shared and Distinct Functions of Type I and Type III Interferons. Immunity 50, 907–923, doi:10.1016/j.immuni.2019.03.025 (2019).

20 Monk, P. D. et al. Safety and efficacy of inhaled nebulised interferon beta-1a (SNG001) for treatment of SARS-CoV-2 infection: a randomised, double-blind, placebo-controlled, phase 2 trial. Lancet Respir Med 9, 196–206, doi:10.1016/S2213-2600(20)30511-7 (2021).

21 Zhou, Q. et al. Interferon-alpha2b Treatment for COVID-19 Is Associated with Improvements in Lung Abnormalities. Viruses 13, doi:10.3390/v13010044 (2020).

22 Hung, I. F. et al. Triple combination of interferon beta-1b, lopinavir-ritonavir, and ribavirin in the treatment of patients admitted to hospital with COVID-19: an open-label, randomised, phase 2 trial. Lancet 395, 1695–1704, doi:10.1016/S0140-6736(20)31042-4 (2020).

23 Lokugamage, K. G. et al. Type I Interferon Susceptibility Distinguishes SARS-CoV-2 from SARS-CoV. J Virol 94, doi:10.1128/JVI.01410-20 (2020).

24 Pfaender, S. et al. LY6E impairs coronavirus fusion and confers immune control of viral disease. Nat Microbiol 5, 1330–1339, doi:10.1038/s41564-020-0769-y (2020).

25 Lei, X. et al. Activation and evasion of type I interferon responses by SARS-CoV-2. Nat Commun 11, 3810, doi:10.1038/s41467-020-17665-9 (2020).

26 Miorin, L. et al. SARS-CoV-2 Orf6 hijacks Nup98 to block STAT nuclear import and antagonize interferon signaling. Proc Natl Acad Sci U S A 117, 28344–28354, doi:10.1073/pnas.2016650117 (2020).

27 Kopecky-Bromberg, S. A., Martinez-Sobrido, L., Frieman, M., Baric, R. A. & Palese, P. Severe acute respiratory syndrome coronavirus open reading frame (ORF) 3b, ORF 6, and nucleocapsid proteins function as interferon antagonists. J Virol 81, 548–557, doi:10.1128/JVI.01782-06 (2007).

28 Nienhold, R. et al. Two distinct immunopathological profiles in autopsy lungs of COVID-19. Nat Commun 11, 5086, doi:10.1038/s41467-020-18854-2 (2020).

29 Sleijfer, S., Bannink, M., Van Gool, A. R., Kruit, W. H. & Stoter, G. Side effects of interferon-alpha therapy. Pharm World Sci 27, 423–431, doi:10.1007/s11096-005-1319-7 (2005).

30 Stanifer, M. L. et al. Critical Role of Type III Interferon in Controlling SARS-CoV-2 Infection in Human Intestinal Epithelial Cells. Cell Rep 32, 107863, doi:10.1016/j.celrep.2020.107863 (2020).

31 Vanderheiden, A. et al. Type I and Type III Interferons Restrict SARS-CoV-2 Infection of Human Airway Epithelial Cultures. J Virol 94, doi:10.1128/JVI.00985-20 (2020).

32 Scholer, L. et al. A Novel In-Cell ELISA Assay Allows Rapid and Automated Quantification of SARS-CoV-2 to Analyze Neutralizing Antibodies and Antiviral Compounds. Front Immunol 11, 573526, doi:10.3389/fimmu.2020.573526 (2020).

33 Desmyter, J., Melnick, J. L. & Rawls, W. E. Defectiveness of interferon production and of rubella virus interference in a line of African green monkey kidney cells (Vero). J Virol 2, 955–961, doi:10.1128/JVI.2.10.955-961.1968 (1968).

34 Jonsdottir, H. R. & Dijkman, R. Coronaviruses and the human airway: a universal system for virus-host interaction studies. Virology journal 13, 24, doi:10.1186/s12985-016-0479-5 (2016).

35 Heinen, N., Klohn, M., Steinmann, E. & Pfaender, S. In Vitro Lung Models and Their Application to Study SARS-CoV-2 Pathogenesis and Disease. Viruses 13, doi:10.3390/v13050792 (2021).

36 V’Kovski, P. et al. Disparate temperature-dependent virus-host dynamics for SARS-CoV-2 and SARS-CoV in the human respiratory epithelium. PLoS Biol 19, e3001158, doi:10.1371/journal.pbio.3001158 (2021).

37 Megger, D. A., Philipp, J., Le-Trilling, V. T. K., Sitek, B. & Trilling, M. Deciphering of the Human Interferon-Regulated Proteome by Mass Spectrometry-Based Quantitative Analysis Reveals Extent and Dynamics of Protein Induction and Repression. Front Immunol 8, 1139, doi:10.3389/fimmu.2017.01139 (2017).

38 Trilling, M. et al. Deciphering the modulation of gene expression by type I and II interferons combining 4sU-tagging, translational arrest and in silico promoter analysis. Nucleic Acids Res 41, 8107–8125, doi:10.1093/nar/gkt589 (2013).

39 Chen, P. et al. SARS-CoV-2 Neutralizing Antibody LY-CoV555 in Outpatients with Covid-19. N Engl J Med 384, 229–237, doi:10.1056/NEJMoa2029849 (2021).

40 Gottlieb, R. L. et al. Effect of Bamlanivimab as Monotherapy or in Combination With Etesevimab on Viral Load in Patients With Mild to Moderate COVID-19: A Randomized Clinical Trial. Jama 325, 632–644, doi:10.1001/jama.2021.0202 (2021).

41 Consortium, W. H. O. S. T. et al. Repurposed Antiviral Drugs for Covid-19 - Interim WHO Solidarity Trial Results. N Engl J Med 384, 497–511, doi:10.1056/NEJMoa2023184 (2021).

42 Hadjadj, J. et al. Impaired type I interferon activity and inflammatory responses in severe COVID-19 patients. Science 369, 718–724, doi:10.1126/science.abc6027 (2020).

43 King, C. & Sprent, J. Dual Nature of Type I Interferons in SARS-CoV-2-Induced Inflammation. Trends Immunol 42, 312–322, doi:10.1016/j.it.2021.02.003 (2021).

44 Bastard, P. et al. Autoantibodies against type I IFNs in patients with life-threatening COVID-19. Science 370, doi:10.1126/science.abd4585 (2020).

45 Zhang, Q. et al. Inborn errors of type I IFN immunity in patients with life-threatening COVID-19. Science 370, doi:10.1126/science.abd4570 (2020).

46 Levy, R. et al. IFN-alpha2a Therapy in Two Patients with Inborn Errors of TLR3 and IRF3 Infected with SARS-CoV-2. J Clin Immunol 41, 26–27, doi:10.1007/s10875-020-00933-0 (2021).

47 Troya, J. et al. Neutralizing Autoantibodies to Type I IFNs in >10% of Patients with Severe COVID-19 Pneumonia Hospitalized in Madrid, Spain. J Clin Immunol, doi:10.1007/s10875-021-01036-0 (2021).

48 Sa Ribero, M., Jouvenet, N., Dreux, M. & Nisole, S. Interplay between SARS-CoV-2 and the type I interferon response. PLoS Pathog 16, e1008737, doi:10.1371/journal.ppat.1008737 (2020).

49 Guo, K. et al. Qualitative Differences Between the IFNalpha subtypes and IFNbeta Influence Chronic Mucosal HIV-1 Pathogenesis. PLoS Pathog 16, e1008986, doi:10.1371/journal.ppat.1008986 (2020).

50 Schlaepfer, E. et al. Dose-Dependent Differences in HIV Inhibition by Different Interferon Alpha Subtypes While Having Overall Similar Biologic Effects. mSphere 4, doi:10.1128/mSphere.00637-18 (2019).

51 Rusinova, I. et al. Interferome v2.0: an updated database of annotated interferon-regulated genes. Nucleic Acids Res 41, D1040–1046, doi:10.1093/nar/gks1215 (2013).

52 Meng, Z. et al. The effect of recombinant human interferon alpha nasal drops to prevent COVID-19 pneumonia for medical staff in an epidemic area. Curr Top Med Chem, doi:10.2174/1568026621666210429083050 (2021).

53 Hoagland, D. A. et al. Leveraging the antiviral type I interferon system as a first line of defense against SARS-CoV-2 pathogenicity. Immunity 54, 557–570 e555, doi:10.1016/j.immuni.2021.01.017 (2021).

54 Hiemstra, P. S., McCray, P. B., Jr. & Bals, R. The innate immune function of airway epithelial cells in inflammatory lung disease. Eur Respir J 45, 1150–1162, doi:10.1183/09031936.00141514 (2015).

55 Li, X., Wang, L., Nunes, D. P., Troxler, R. F. & Offner, G. D. Pro-inflammatory cytokines up-regulate MUC1 gene expression in oral epithelial cells. J Dent Res 82, 883–887, doi:10.1177/154405910308201107 (2003).

56 Chatterjee, M., van Putten, J. P. M. & Strijbis, K. Defensive Properties of Mucin Glycoproteins during Respiratory Infections-Relevance for SARS-CoV-2. mBio 11, doi:10.1128/mBio.02374-20 (2020).

57 Sungnak, W. et al. SARS-CoV-2 entry factors are highly expressed in nasal epithelial cells together with innate immune genes. Nat Med 26, 681–687, doi:10.1038/s41591-020-0868-6 (2020).

58 Zou, X. et al. Single-cell RNA-seq data analysis on the receptor ACE2 expression reveals the potential risk of different human organs vulnerable to 2019-nCoV infection. Front Med 14, 185–192, doi:10.1007/s11684-020-0754-0 (2020).

59 Lu, W. et al. Elevated MUC1 and MUC5AC mucin protein levels in airway mucus of critical ill COVID-19 patients. J Med Virol 93, 582–584, doi:10.1002/jmv.26406 (2021).

60 Jonsdottir, H. R. & Dijkman, R. Characterization of human coronaviruses on well-differentiated human airway epithelial cell cultures. Methods Mol Biol 1282, 73–87, doi:10.1007/978-1-4939-2438-7_8 (2015).

61 Livak, K. J. & Schmittgen, T. D. Analysis of relative gene expression data using real-time quantitative PCR and the 2(−Delta Delta C(T)) Method. Methods 25, 402–408, doi:10.1006/meth.2001.1262 (2001).

62 Hu, J., Ge, H., Newman, M. & Liu, K. OSA: a fast and accurate alignment tool for RNA-Seq. Bioinformatics 28, 1933–1934, doi:10.1093/bioinformatics/bts294 (2012).

63 Love, M. I., Huber, W. & Anders, S. Moderated estimation of fold change and dispersion for RNA-seq data with DESeq2. Genome Biol 15, 550, doi:10.1186/s13059-014-0550-8 (2014).

64 Kramer, A., Green, J., Pollard, J., Jr. & Tugendreich, S. Causal analysis approaches in Ingenuity Pathway Analysis. Bioinformatics 30, 523–530, doi:10.1093/bioinformatics/btt703 (2014).

65 Ianevski, A., He, L., Aittokallio, T. & Tang, J. SynergyFinder: a web application for analyzing drug combination dose-response matrix data. Bioinformatics 33, 2413–2415, doi:10.1093/bioinformatics/btx162 (2017).

